# Dynamic analysis of sugar metabolism reveals the mechanisms of action of synthetic sugar analogs

**DOI:** 10.1101/2020.09.15.288712

**Authors:** Monique van Scherpenzeel, Federica Conte, Christian Büll, Angel Ashikov, Esther Hermans, Anke Willems, Walinka van Tol, Else Kragt, Ed E. Moret, Torben Heise, Jeroen D. Langereis, Emiel Rossing, Michael Zimmermann, M. Estela Rubio-Gozalbo, Marien I. de Jonge, Gosse J. Adema, Nicola Zamboni, Thomas Boltje, Dirk J. Lefeber

## Abstract

Synthetic sugar analogs are widely applied in metabolic oligosaccharide engineering (MOE) and as novel drugs to interfere with glycoconjugate biosynthesis. However, mechanistic insights on their exact metabolism in the cell and over time are mostly lacking. We developed sensitive ion-pair UHPLC-QqQ mass spectrometry methodology for analysis of sugar metabolites in organisms and in model cells and identified novel low abundant nucleotide sugars in human cells, such as ADP-glucose and UDP-arabinose, and CMP-sialic acid (CMP-NeuNAc) in Drosophila. Dynamic tracing of propargyloxycarbonyl (Poc) labeled analogs, commonly used for MOE, revealed that ManNPoc is metabolized to both CMP-NeuNPoc and UDP-GlcNPoc. Finally, combined treatment of B16-F10 melanoma cells with antitumor compound 3F_ax_-NeuNAc and ^13^C-labeled GlcNAc revealed that endogenous CMP-NeuNAc levels started to decrease before a subsequent decrease of ManNAc 6-phosphate was observed. This implicates 3F_ax_-NeuNAc first acts as a substrate for cytosolic CMP-sialic acid synthetase and subsequently its product CMP-3F_ax_-NeuNAc functions as a feed-back inhibitor for UDP-GlcNAc 2-epimerase/N-acetylmannosamine kinase. Thus, dynamic analysis of sugar metabolites provides key insights into the time-dependent metabolism of synthetic sugars, which is important for the rational design of analogs with optimized effects.

## Introduction

Monosaccharides serve as important metabolic intermediates for the production of energy and glycoconjugates across all domains of life. During sugar metabolism, monosaccharides are converted to their corresponding nucleotide sugars in the cellular cytoplasm in a complex metabolic network involving numerous metabolic intermediates and more than 40 currently known enzymes in mammals (Figure 1 a). Nucleotide sugars serve as building block for glycoproteins, glycolipids and glycosylphosphatidylinositol anchors. These complex glycoconjugates are involved in important biological processes such as cellular communication and signaling and have been associated with numerous human diseases, both acquired^[1–4]^ and genetic.^[5]^ Disruptions in sugar metabolism have been linked to various diseases such as cancer,^[6]^ neurodegenerative^[7]^ and infectious^[8]^ disease, some of them being treatable via dietary intervention in sugar metabolism like galactose supplementation in PGM1 myopathy.^[9]^ Its ease of accessibility has made sugar metabolism an attractive target to design synthetic sugar analogs to restore abnormal glycosylation and to study the biological role of glycoconjugates in cells and organisms by metabolic oligosaccharide engineering (MOE).

**Figure 1.**
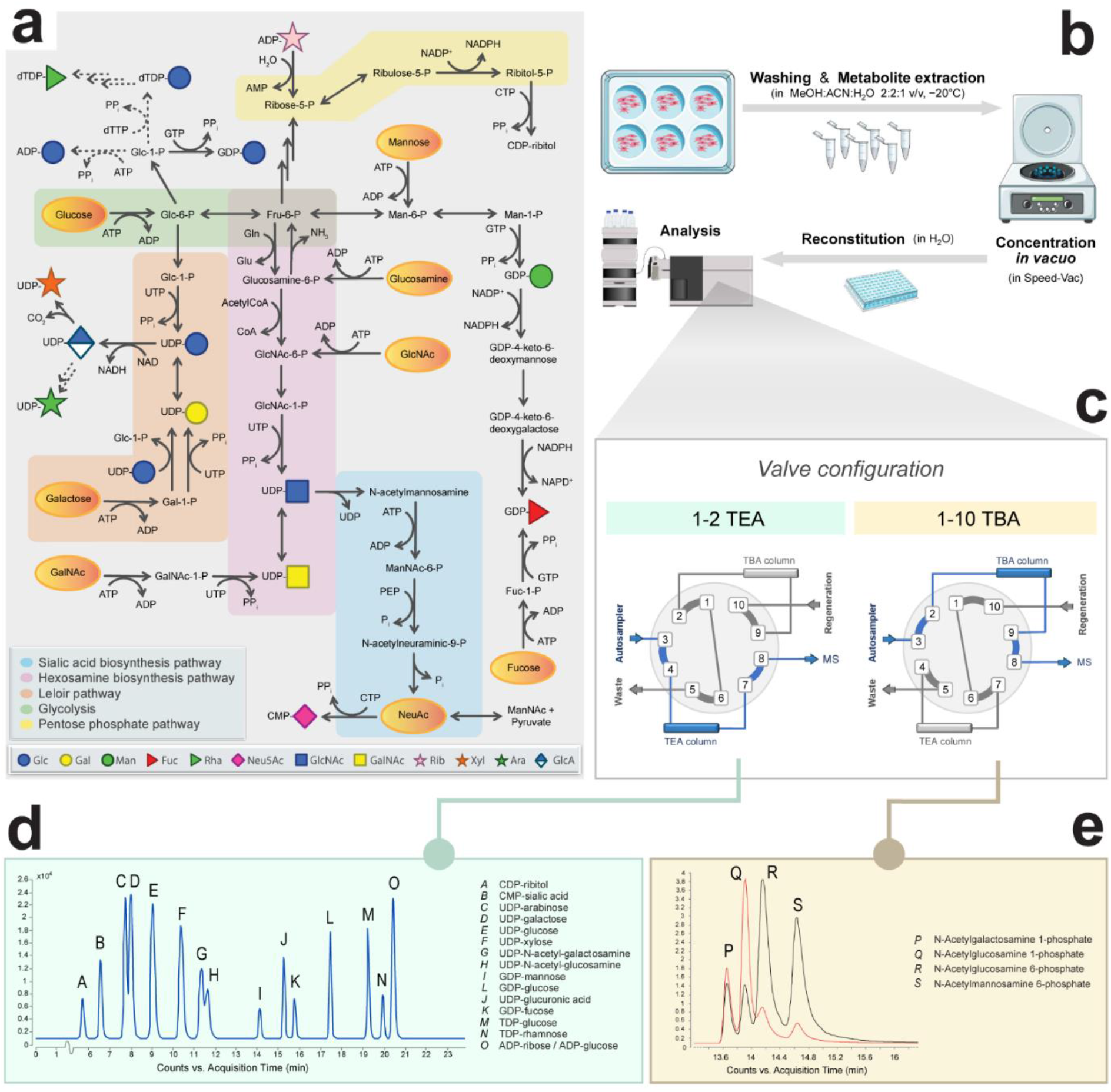
Sugar metabolism as integral part of intracellular metabolism. a) Overview of the metabolic pathways involved in the generation of nucleotide sugars and their intermediate sugar phosphates in humans. Dashed arrows indicate pathways that remain to be identified in humans. Yellow ovals indicate salvage pathways where supplemental sugars can enter the metabolic pathways. Abbreviations: GLN, glutamine; GLU, glutamate; PEP, phosphoenolpyruvate; PPi, pyrophosphate; Pi, inorganic phosphate; −P, phosphate; Glc, glucose; Gal, galactose; Fuc, fucose; Fru, fructose; Man, mannose; NeuNAc, *N*-acetylneuraminic acid; GlcNAc, *N*-acetylglucosamine; GalNAc, *N*-acetylgalactosamine; ManNAc, *N*-acetylmannosamine; AMP/ADP/ATP, adenosine 5’-mono-/di-/triphosphate; CDP/CTP, cytidine 5’-di-/triphosphate; GDP/GTP, guanosine 5’-di-/triphosphate; dTDP/dTTP, deoxythymidine di-/triphosphate; UDP/UTP, uridine 5’-di-/triphosphate; CoA, coenzyme A; NAD(H), nicotinamide adenine dinucleotide; NADP(H), nicotinamide adenine dinucleotide phosphate. b) Sample preparation consists of 6-well cell cultures, followed by extraction of intracellular metabolites for subsequent UHPLC-QqQ analysis. c) The samples can be injected twice via a versatile and automatic switching valve to switch between two different ion pair mobile phases. d) Typical chromatogram of 16 nucleotide sugar standards. e) A selection of *N*-acetylhexosamine phosphate sugars is shown. The red line illustrates the MRM transition 300.0 -> 79.0 m/z, the black line MRM transition 300.0 -> 97.0 m/z.

In MOE, monosaccharides with different chemical groups, such as azide, alkynyl or propargyloxycarbonyl (Poc), enter sugar metabolism via salvage pathways, and are subsequently integrated into glycoconjugates by glycosyltransferases in the ER and Golgi apparatus. These chemical groups can then be indirectly visualized after bioorthogonal chemistry by fluorescent detection or mass spectrometry.^[10–13]^ Although widely applied, only few studies report on the different substrate specificities of enzymes in sugar metabolism for specific chemical reporter groups. For example, alkyne functionalized GlcNAc (GlcNAlk) was found to be more specific for labeling of O-GlcNAcylated proteins than azide functionalized GlcNAc (GlcNAz) since it was not accepted as substrate by the cytosolic enzyme GALE.^[14]^ In addition, specific sialic acid analogs were not accepted as substrate for CMP-sialic acid synthetase (CMAS).^[15]^

Synthetic sugars are also developed as inhibitors of sugar metabolic pathways with the aim to develop novel therapeutics,^[16]^ such as antimicrobial^[17,18]^ and anticancer^[19–21]^ drugs. Similarly, they are used to study the role of glycoconjugates in biological systems. For instance, 6-Diazo-5-oxo-L-norleucine is used to inhibit the synthesis of UDP-GlcNAc. However, being a glutamine analog, it also inhibits other glutamine-dependent enzymes such as glutaminase.^[22]^ Similarly, 2-deoxyglycose, originally designed as inhibitor of glycolysis, has been shown to exert some of its biological effects by inhibiting protein N-glycosylation via the GDP-mannose pathway.^[23]^ Moreover, novel nucleotide sugars have recently been identified in human cells and tissues, including UDP-mannose^[24]^ and CDP-ribitol,^[25]^ the latter being targeted to develop antimycobacterial drugs^[17]^ as it was supposed to be absent in human. Thus, care should be taken how to interpret the results of studies using synthetic sugars. Together with recent evidence for the existence of additional enzymes in sugar metabolism, the molecular mechanisms and interplay between pathways appear far more complex than anticipated.

To further optimize the effectiveness and specificity of synthetic sugar analogs, detailed knowledge on their metabolism is becoming important. Studies on individual enzymes have provided information on enzyme specificities in different species, however, a more holistic view on cellular metabolism is highly warranted to investigate the dynamics of incorporation of synthetic sugar analogs in metabolic pathways and their side effects. Separation of nucleotide sugars and nucleotides has classically been performed by HPLC with triethylamine (TEA) ion-pairing buffer.^[26,27]^ This was later combined with ion-trap^[28]^ or triple quad^[29]^ mass spectrometry to increase sensitivity and scale down cell culture to 6-well plates. Dedicated methods for analysis of the sugar phosphate intermediates are lacking. However, a number of pentose and hexose phosphates can be analyzed using analytical methods for central carbon metabolism, including glycolysis and pentose-phosphate pathway (PPP), based on chromatographic separation using tributylamine (TBA) ion-pairing buffer.^[30]^ Here, we report ultrahigh performance liquid chromatography – triple quadrupole mass spectrometry (UHPLC-QqQ) methodology using both TEA and TBA buffers for detailed analysis of the metabolism of synthetic sugars. We first profiled an extended set of nucleotide sugars in 13 commonly used model cell lines and organisms and identified the presence of novel low abundant nucleotide sugars. Secondly, we studied the dynamics of commonly used Poc derivatives for MOE and could confirm the metabolism of ManNPoc towards UDP-GlcNPoc. Lastly, we combined the two ion-pairing methods to investigate the mechanism of action of the potent anti-tumor sugar analog 3F_ax_-NeuNAc, by tracing isotopically labeled ^13^C-labeled GlcNAc in CMP-NeuNAc metabolism. In addition to a depletion of CMP-NeuNAc and production of CMP-3F_ax_-NeuNAc, a time-dependent strong decrease was identified of *N*-acetyl-mannosamine 6-phosphate (ManNAc-6P), indicating feed-back inhibition of the produced CMP-3F_ax_-NeuNAc on UDP-GlcNAc 2-epimerase/N-acetylmannosamine kinase (GNE).

## Results and Discussion

### UHPLC-QqQ mass spectrometry for analysis of sugar metabolites in model cell lines and organisms

With the aim to establish a single method for analysis of all sugar metabolites, we elaborated on a reverse phase TBA ion-pairing UHPLC-QqQ mass spectrometry method for analysis of polar and anionic metabolites in central carbon metabolism.^[30]^ We selected a set of metabolites representative of glycolysis and the pentose phosphate pathway (PPP), also in view of their known correlation with sugar metabolism, such as the selective shutdown of the GDP-mannose pathway upon glucose deprivation,^[31]^ and added pentose- and hexose-phosphates and *N*-acetylhexosamine phosphates by analysis of commercial standards (Figure 1 d). Nucleotide sugars could be analyzed as well, however, as they eluted at the end of the gradient as broad peaks, separation of isobaric nucleotide sugars was impossible. Therefore, dedicated analysis of nucleotide sugars was performed by separation of 18 nucleotide sugars with 20 mM TEA buffer,^[27]^ resulting in separation of isobaric metabolites, like UDP-glucose and UDP-galactose, and of UDP-GalNAc and UDP-GlcNAc (Figure 1 d). MRM transitions were obtained via direct infusion of a set of 14 commercially available standards (Table S1) that were also used to optimize ion source settings. Linearity, lower limit of detection (LOD) and lower limit of quantification (LOQ) were determined as validation in control primary human fibroblasts (Table S2). In brief, low nanomolar concentrations could be detected, the linearity spanned three orders of magnitude with R^2^>0.99, carry-over was <0.1% for all nucleotide sugars and inter- and intraday coefficients of variation (n=10) were below 10% for all compounds. To enable analysis of both nucleotide sugars and sugar-phosphate intermediates, a switching valve was introduced and two similar columns to run batches of TEA and TBA analyses from single samples (Figure 1 b and c).

As a first step towards diverse applications, we profiled nucleotide sugars in nine commonly studied model cell lines and organisms (Figure 2 and Table S4). We added UDP-ManNAc and TDP-rhamnose in view of their importance in bacteria and CMP-NeuGc in view of its importance in mouse cell lines (for conditions, see Table S1). Overall, the three human cell lines showed a similar relative distribution with some differences for example in CMP-NeuNAc and UDP-GlcA. For human HAP1 cells, steady state levels were not significantly influenced by cell culture conditions (Figure S1 and Table S4). Interestingly, low relative levels of ADP-, TDP- and GDP-glucose could be identified, previously only described in non-human species, such as ADP-glucose in plants. In addition, traces of UDP-arabinose were detected, previously identified in plants but also in CHO cells.^[32]^ For C2C12 mouse myoblasts, a strong change in the relative distribution of nucleotide sugars was observed during differentiation towards myotubes with a strong increase in the levels of CDP-ribitol and UDP-GlcNAc and a relative decrease in UDP-glucose. CDP-ribitol was recently identified in human muscle tissue and is important for the glycosylation of the muscle membrane protein alpha-dystroglycan^[33]^ and might therefore be upregulated during muscle cell differentiation.

**Figure 2.**
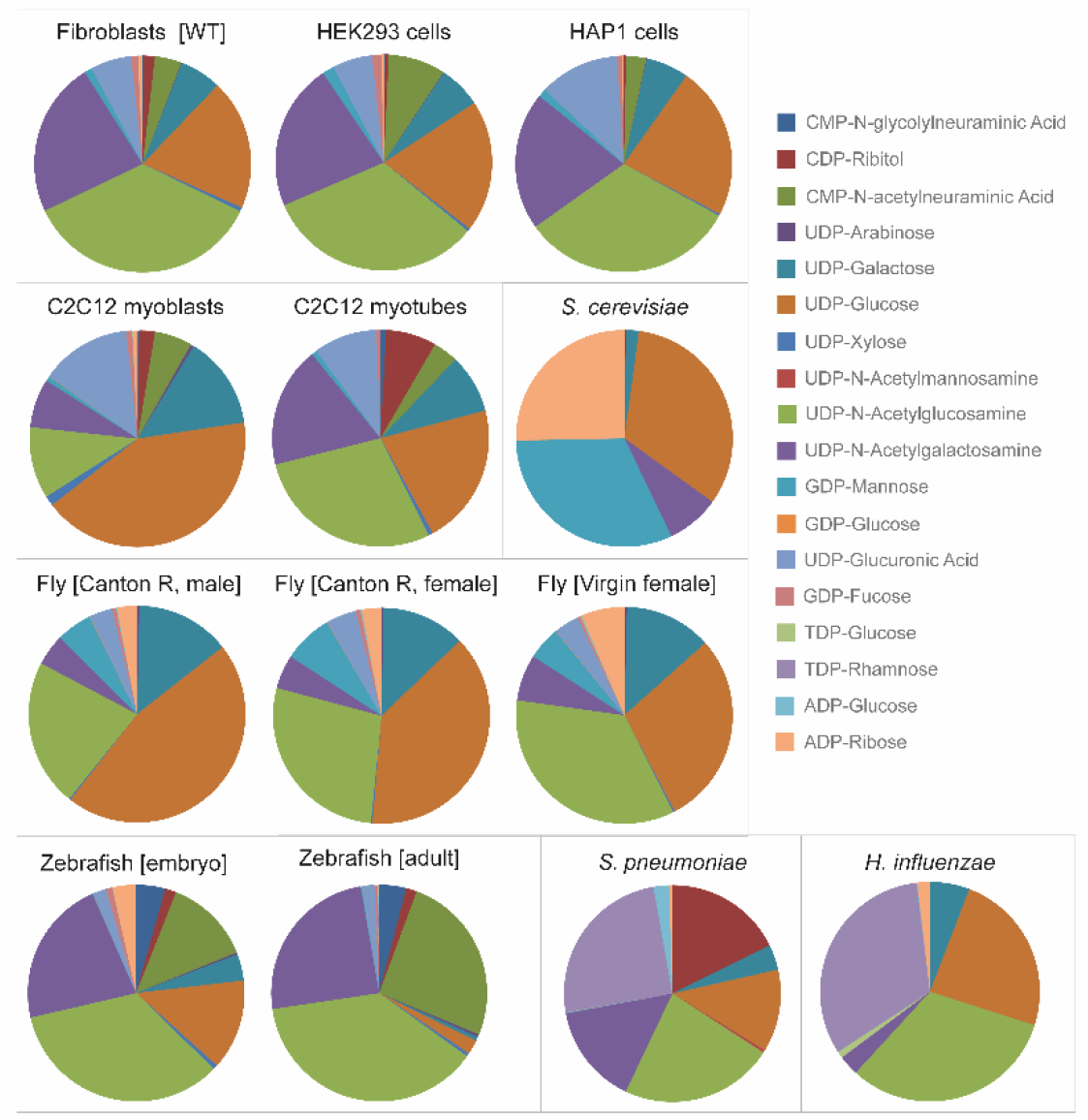
Nucleotide sugar profiles of 13 common model organisms and cell lines. Peak intensities of 18 nucleotide sugars (Table S1) were normalized by the sum of all nucleotide sugars in a given sample and expressed as relative percentage in pie charts. Organisms are shown at the bottom and cell lines at the top. Cell models: human dermal fibroblast (wild-type, WT), human embryonic kidney HEK293 cells, human haploid HAP1 cells, murine C2C12 myoblasts, murine C2C12 myotubes. Model organisms: *Saccharomyces cerevisiae* (yeast), *Drosophila melanogaster* (fly), *Danio rerio* (zebrafish), *Streptococcus pneumoniae* (gram-positive bacteria), *Haemophilus influenzae* (gram-negative bacteria).

Each organism showed its own typical nucleotide sugar profile and could be separated by hierarchical clustering analysis (Figure S2). For example, TDP-rhamnose was detected in *Sreptococcus pneumonia* and *Haemophilus influenzae* and UDP-ManNAc and CDP-ribitol in *H. influenzae*, which are essential nucleotide sugars for synthesis of capsular polysaccharides in these bacteria.^[34]^ GDP-mannose levels were most dominant in yeast, as expected because of the high need for mannoprotein synthesis, but were also found at relatively high levels in *Drosophila melanogaster*. In addition, we could identify low relative levels of CMP-NeuNAc in *D. melanogaster*. Although the metabolic pathway for CMP-NeuNAc synthesis has not been broadly investigated in *D. melanogaster*, our finding is in agreement with the presence of sialic acid in *D. melanogaster* N-glycans^[35]^ and the intracellular localization of CMP-NeuNAc synthetase, and provides a starting point for MOE.

### Metabolic incorporation of Poc labeled sugars in sialic acid biosynthesis

Over the last decade, MOE has developed as a field to label glycoconjugates by use of synthetic sugars for visualization of cellular glycans or for isolation and subsequent glycomics analysis. A plethora of differently modified monosaccharides is being used for incubation of cells or model organisms with varying labeling efficiencies, possibly related to differences in their metabolism. In previous studies, ManNPoc was shown to be incorporated into glycoconjugates as sialic acid derivative. Its incorporation in cellular glycoconjugates after complete inhibition of sialic acid biosynthesis^[36]^ suggested an additional metabolic fate. To study this, we incubated human fibroblasts with Ac4ManNPoc and Ac5NeuNPoc and programmed MRM transitions of UDP-GlcNPoc and CMP-NeuNPoc, based on the theoretical mass and fragments of non-derivatized nucleotide sugars (Figure S3 a, Figure 3 d, Table S1). After 48 hours of incubation with NeuNPoc, the Poc label exclusively appeared in CMP-NeuNPoc (for 32%). When ManNPoc was administered to cells, 18% labeling was observed in CMP-NeuNPoc, while additionally UDP-HexNPoc was formed for 0.6% as compared to endogenous UDP-GlcNAc (Figure 3 a and Figure S3 b). Incubation with ManNPoc or NeuNPoc slightly reduced the levels of endogenous CMP-NeuNAc, while the relative levels of other nucleotide sugars remained unaffected (Table S5).

**Figure 3.**
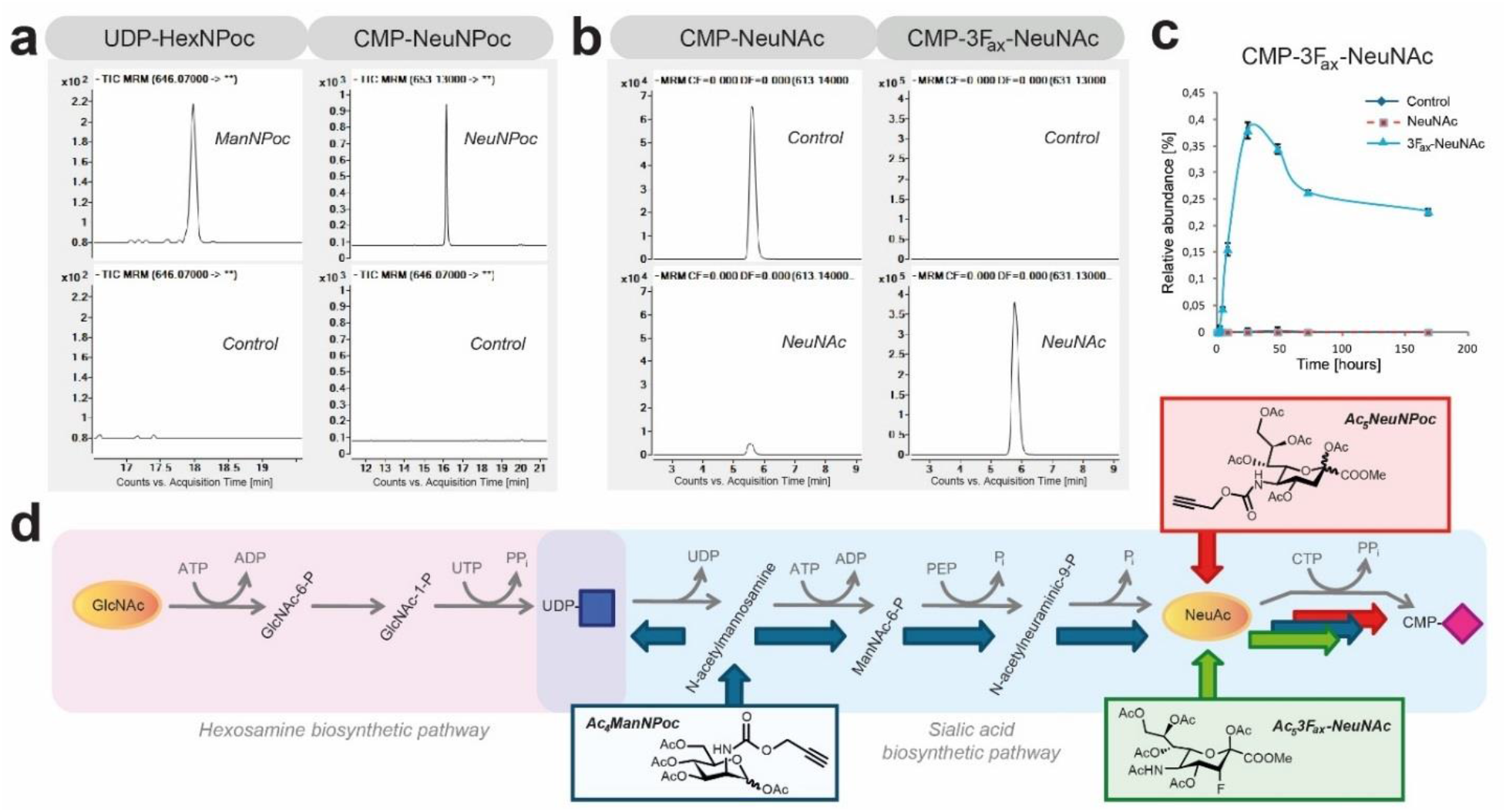
Incorporation of chemical reporter groups in nucleotide sugars. a) Incubation of fibroblasts for 48 h with ManNPoc results in metabolic labeling of both UDP-GlcNPoc and CMP-NeuNPoc, while these nucleotide sugars were absent from controls. b) Formation of CMP-3F_ax_-NeuNAc by feeding B16-F10 cells with 3F_ax_-NeuNAc for 24 h and not by control incubations. c) Time-dependent formation of CMP-3F_ax_-NeuNAc. d) Shown are the *de novo* CMP-NeuNAc biosynthesis pathway and the salvage pathway of GlcNAc. Indicated in blue arrows is the metabolic route of incorporation of ManNPoc, in red of NeuNPoc and in green of 3F_ax_-NeuNAc.

### Dynamic tracing of *de novo* CMP-NeuNAc synthesis with reveals the cytosolic mechanism of action of sialylation inhibitor 3F_ax_-NeuNAc

As a second chemical glycobiology application to show the importance of analyzing intermediary sugar phosphates, we applied our methodology to study the mechanism of a synthetic inhibitor for sialylation. Global hypersialylation in cancer, as observed for over four decades,^[37,38]^ can be inhibited by administration of a fluorinated sialic acid analogue, 3F_ax_-NeuNAc (Figure S3 a), resulting in a reduction in tumor growth in mice.^[39]^ Its mechanism of action is proposed to be based on the inhibition of Golgi localized sialyltransferases as confirmed by *in vitro* enzyme assays.^[40]^ *In vivo*, neosynthesized CMP-3F_ax_-NeuNAc should be subsequently transported into the Golgi, likely via the CMP-NeuNAc transporter SLC35A1, to inhibit sialyltransferase isoenzymes. To study the metabolism of 3F_ax_-NeuNAc in more detail, we combined our TEA and TBA analytical assays to profile both nucleotide sugars and sugar phosphates intermediates of the hexosamine and sialic acid pathways. We first incubated B16-F10 melanoma cells with peracetylated 3F_ax_-NeuNAc and evaluated its metabolism over time. Following a single pulse, time-dependent formation was observed of CMP-3F_ax_-NeuNAc, with a maximum accumulation at 24 hours which decreased to about 60-70% of maximum levels and remained stably detectable up to at least seven days (Figure 3 c). The effect on endogenous metabolites was monitored for 48 hours. A reduction of CMP-NeuNAc appeared after about 4 hours, with a further reduction to near depletion after 24 hours (Figure 4 a). In addition, analysis of the intermediate sugar phosphates revealed a marked reduction of ManNAc-6P levels, becoming apparent after 24 hours when maximum levels of synthesized CMP-3F_ax_-NeuNAc were reached. No effects on the levels of other endogenous metabolites were observed, neither were the levels of ManNAc-6P influenced by incubations with PBS or peracetylated NeuNAc as controls. The relative levels of other nucleotide sugars were slightly reduced, but this is explained by the high levels of produced CMP-3F_ax_-NeuNAc (Table S6). These results suggest that in first instance, 3F_ax_-NeuNAc is metabolized to its corresponding CMP derivative, while it also competes with endogenous NeuNAc for CMP-NeuNAc synthesis, thereby initiating the decrease in CMP-NeuNAc levels. Subsequently, *de novo* CMP-NeuNAc synthesis by GNE is inhibited by feed-back inhibition by CMP-3F_ax_-NeuNAc.

**Figure 4.**
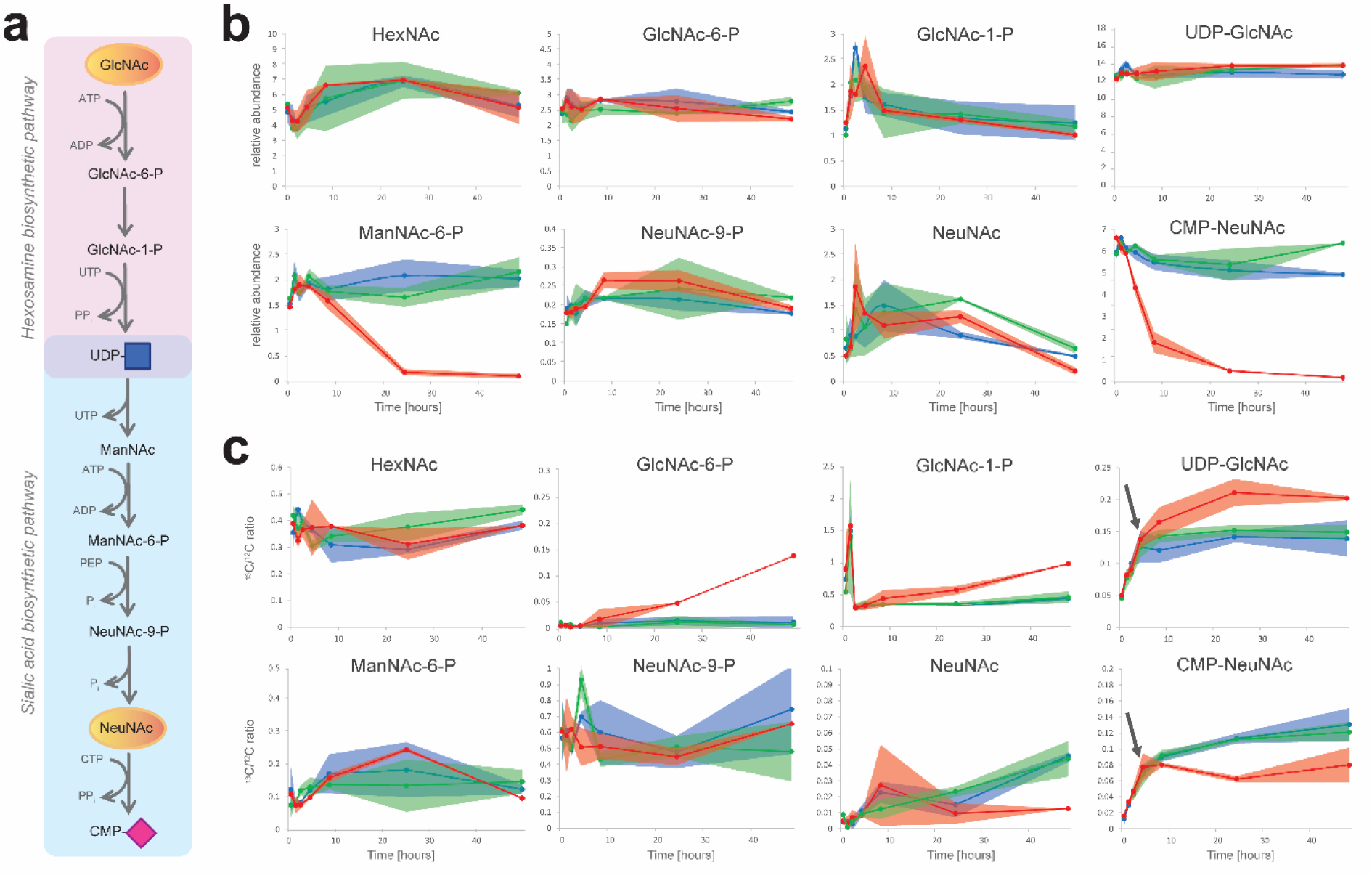
Effect of 3F_ax_-NeuNAc on sialic acid metabolism. a) Position of the presented metabolites in the metabolic pathway. b) B16-F10 mouse myeloma cells were incubated with 100 microM 3F_ax_-NeuNAc for 5 min, 1, 2, 4, 8, 24, and 48 hours (red line). Incubations with PBS (blue line) and Ac5NeuNAc (green line) for the same time points were performed as control. Normalized relative levels of sugar metabolites are presented. c) B16-F10 mouse myeloma cells were incubated with 100 microM 3F_ax_-NeuNAc for the same time points as in b) in the presence of 1 mM [UL-^13^C6]-GlcNAc (red line). Incubations with PBS (blue line) and Ac_5_NeuNAc (green line) for the same time points, both in the presence of 1 mM [UL-^13^C6]-GlcNAc, were performed as control.

To obtain further insight into the *de novo* biosynthesis pathway of CMP-NeuNAc in the presence of 3F_ax_-NeuNAc and the involvement of GNE, we performed metabolite tracing of isotopically labeled *N*-acetyl-D-[UL-^13^C6]-glucosamine ^13^C6-GlcNAc) in the presence of 3F_ax_-NeuNAc. Theoretical MRMs for ^13^C6-labeled intermediates of the pathway (Figure 4 a) were programmed and data expressed as ^13^C/^12^C-labeling ratio (Figure 4 c). In control incubations, ^13^C6-GlcNAc was metabolized to UDP-^13^C6-GlcNAc and CMP-^13^C6-NeuNAc in a time-dependent manner reaching a plateau after about 8 hours. In the presence of 3F_ax_-NeuNAc, a relative accumulation of UDP-^13^C6-GlcNAc started to appear between 4 and 8 hours, at the same time when metabolism towards CMP-^13^C6-NeuNAc started to decrease. This is suggestive for an inhibition of GNE enzyme activity, as accumulating UDP-GlcNAc is less efficiently converted to ManNAc-6P. More or less in the same time frame, the relative levels of ^13^C6-GlcNAc-6P and ^13^C6-GlcNAc-1P increased as compared to control incubations with PBS and acetylated NeuNAc, indicating a block in the pathway beyond UDP-GlcNAc. Taken together, our data on ^13^C-GlcNAc tracing confirm the allosteric inhibition of GNE by CMP-3F_ax_-NeuNAc, which is formed in sufficient amounts after about 4-8 hours to affect *de novo* CMP-NeuNAc synthesis.

We next compared the binding of 3F_ax_-NeuNAc and NeuNAc to CMAS and of their CMP derivatives to GNE by molecular modeling. NeuNAc was redocked and F-NeuNAc was docked in the A site of the AB dimer of CMAS,^[41]^ using the X-ray structure of murine CMAS as model for the active site of the enzyme. Both molecules overlapped and fitted in the narrow active site (Figure 5 a). For GNE, CMP-3F_ax_-NeuNAc and CMP-NeuNAc were docked to a human model of GNE.^[42]^ Both compounds overlapped and bound to the allosteric binding site of GNE (Figure 5 b). Thus, molecular modeling substantiates our findings on the activity of both CMAS and GNE for the fluorinated sialic acid derivatives. Previously, CMP-3F_ax_-NeuNAc was shown to inhibit human sialyltransferase ST6Gal I in *in vitro* enzyme assays.^[40]^ Interestingly, both the axial and the equatorial variant inhibited ST6Gal I *in vitro* with similar IC_50_ (IC_50_-values of 9.5 μM for CMP-3F_ax_-NeuNAc and 4.7 μM for CMP-3F_eq_-NeuNAc). However, 3F_eq_-NeuNAc did not show an inhibitory effect on sialylation when incubated with different cell lines, proposed to be caused by the inability of CMAS to metabolize this compound.^[38]^ With the methodology presented here, studies regarding the cellular metabolism of sugar isomers such as 3F_eq_-NeuNAc now become feasible. A relevant question concerns the kinetics of the different inhibitory effects of 3F_ax_-NeuNAc. Inhibition of cellular sialylation requires a couple of days,^[39]^ while CMP-NeuNAc is already depleted within 24 hours, suggesting fast inhibition of GNE. It is therefore likely that the depletion of cytosolic CMP-NeuNAc in itself contributes to the decrease in cellular sialylation, in addition to the direct inhibition of Golgi sialyltransferases. In further studies, it would therefore be interesting to investigate the transport efficiency of CMP-3F_ax_-NeuNAc into the Golgi via SLC35A1. Moreover, it would be interesting to combine our methodology for tracing of sugar metabolism with recent methodology for tracing of ^13^C labeled monosaccharides into glycoconjugates^[43]^ to dissect the time-dependent contribution of both inhibitory effects on sialylation.

**Figure 5.**
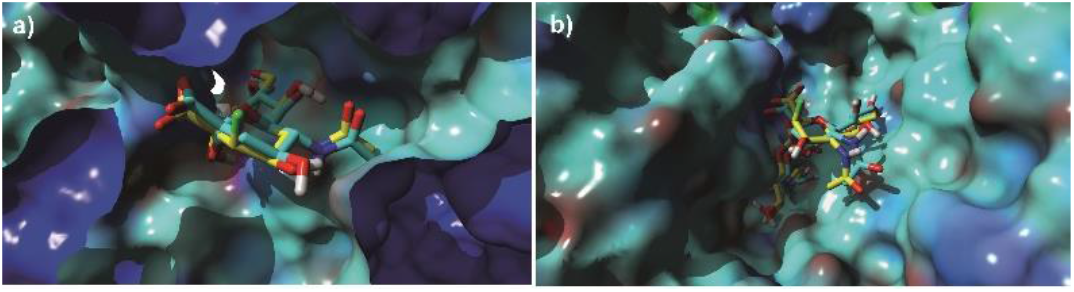
Molecular modeling CMAS and GNE with fluorinated sialic acid analogs. a) Docking of NeuNAc (in blue) and 3F_ax_-NeuNAc (in yellow) in CMAS. b) Docking of CMP-NeuNAc (in blue) and CMP-3F_ax_-NeuNAc (in yellow) in GNE.

## Conclusion

In summary, we established methodology for analysis of nucleotide sugars and sugar phosphates which allows to investigate the effects of synthetic sugar analogs on sugar metabolism in a cellular context, at steady state or by dynamic tracing with isotopically labeled monosaccharides. Separation of isobaric sugar metabolites is challenging and reverse phase chromatography with ion pairing buffers is capable of separating all currently known sugar metabolites. In addition, it allows analysis of additional metabolic pathways in central carbon metabolism, that are interlinked with sugar metabolism. Ideally, a single method would be used for analysis of sugar phosphates and nucleotide sugars, however, TBA ion-pairing buffer did not provide separation of isobaric nucleotide sugars. An additional disadvantage is that the use of a dedicated LC-MS instrument is preferred in view of the contamination with ion-paring agents. Further developments in ion-mobility mass spectrometry might provide a solution to this problem.^[44]^

Our results from steady-state analysis further expand the repertoire of known human nucleotide sugars. For a long time, synthesis of human glycoconjugates was believed to require nine different nucleotide sugars. Identification of ISPD as disease-causing gene^[45]^ resulted in the identification of CDP-ribitol in human cells and tissues^[33]^, while more recently, UDP-mannose was detected in human cell lines and mouse tissues.^[24]^ With increased sensitivity, we could detect additional low abundant nucleotide sugars, ADP-glucose, GDP-glucose, TDP-glucose, and UDP-arabinose. Possibly, these are the byproducts of enzymes that are not 100% specific and form non-classical metabolites at low rates.^[46]^ The relevance of these nucleotide sugars for glycoconjugate biosynthesis is not clear as it is unknown if these could be transported into the Golgi apparatus for glycosylation reactions or they might interfere with physiological glycosylation processes. The identification of TGDS as disease gene^[47]^ suggests the presence of even additional nucleotide sugars in human. Thus, this will impact the development of synthetic sugars as antimicrobial drug by targeting sugar metabolic pathways that now also appear to be present in human, such as the ISPD pathway^[17]^.

Another important factor to consider is the fact that intermediary sugar phosphates can influence more distant sugar metabolic pathways. For example, accumulating fructose 1-phosphate inhibits the enzyme mannose phosphate isomerase in the mannose pathway.^[48]^ Since these effects are not easily predictable, care should be taken when designing synthetic compounds for interfering in sugar metabolism and a broad profiling of sugar metabolites and interconnected metabolic pathways as we presented here is recommended to study the specificity and mechanisms of action of synthetic sugar analogs.

In addition to steady state analysis, dynamic tracing of sugar metabolism with isotopically labeled sugars can provide detailed information on the effect of synthetic sugars on individual enzymes in their cellular context. We provide evidence that ManNPoc can be metabolized towards both UDP-GlcNPoc and CMP-NeuNPoc with different efficiencies, which should be considered for MOE studies. It is highly likely that this will depend on the activities of the different enzymes involved, which will differ between cell types and conditions. For instance, we showed that the levels of specific nucleotide sugars change during muscle cell maturation. While tissue-specific regulation of common metabolic pathways such as glycolysis is widely known, this is hardly studied for sugar metabolism. Nonetheless, evidence for tissue-specific regulation is increasing, for example as recently shown for the muscle-specific role of sialic acid catabolism.^[49]^ Thus the dynamics and directionality of sugar metabolic pathways should be included in the design of new generation synthetic sugars for precise targeting of glycosylation pathways in a more systematic manner.

In conclusion, we provide methodology to study the interconnected networks of central carbon and sugar metabolism to further optimize the design of synthetic sugars for metabolic oligosaccharide engineering and therapeutic strategies.

## Acknowledgements

We would like to thank Dr. Gerrit Hermann from ISOtopic solutions (Austria) for his kind gift of fully ¹³C-labelled yeast extract, and Christa van der Gaast-deJongh for technical support. This study was financially supported by the Netherlands Organization for Scientific Research (VIDI Grant 91713359 to D.J.L), the Prinses Beatrix Spierfonds (Grant W.OR17-15 to D.J.L) and the European Union’s Horizon 2020 research and innovation program under the ERA-NET Cofund action N° 643578 (EUROCDG-2).

