## Supplemental methods and figures for "Dynamic analysis of sugar metabolism reveals the mechanisms of action of synthetic sugar analogs"

### Supporting Information


| | |
|---|---|
| Experimental Section | 3 |
| Supplementary Figure 1: | 8 |
| Supplementary Figure 2: | 9 |
| Supplementary Figure 3: | 10 |
| Supplementary Figure 4: | 11 |
| Supplementary Tables 1-6: see Online Files | |

#### EXPERIMENTAL SECTION

##### Chemicals

Methanol and 2-propanol, both LC-MS grade, the ion pair reagents 3-tributylamine (TBA) and triethylamine-acetic acid (TEA), media, and sample preparation chemicals as well as all commercial standards (Table S1 and S3) at the highest available purity were purchased from Sigma-Aldrich (Switzerland), unless stated otherwise. Nanopure water was obtained with an electric resistance of greater than 10 MΩ from a MilliQ purification unit (Millipore, Bedford, MA). TDP-rhamnose was synthesized upon request by CarboSynth (United Kingdom). All studies are in compliance with local ethical rules.

##### Extraction of polar metabolites from biological samples

###### *Cell lines and incubations*

*Human dermal fibroblasts (primary) and incubation with propargyloxycarbonyl sugars.* Cells were cultured in 6-well plates in M199 medium (PanBioTech), supplemented with 10% fetal calf serum (FCS, Gibco) and 1% Penicilin-Streptamycin (PenStrep, Gibco). After reaching 70-80% confluence, cells were rinsed twice with 75 mM ammonium carbonate (pH=7.4) and plates were immediately frozen in liquid nitrogen and stored at –80°C until metabolite extraction. For analysis of the metabolism of propargyloxycarbonyl derivatized sugars, ManNPoc and SiaNPoc were synthesized as described before.[39] $Ac_4$ManNPoc, $Ac_5$NeuNPoc or $Ac_5$NeuAc as control were diluted in culture medium to a final concentration of 15 µM synthetic sugar in the medium and added to growing cells by refreshing the medium. After 48 hours incubation with synthetic sugar, cells were rinsed twice with 75 mM ammonium carbonate (pH=7.4) and plates were immediately frozen in liquid nitrogen and stored at –80°C until metabolite extraction.

*Human Embryonic Kidney cells (HEK293).* Cells were cultured on 6-well plates in DMEM (Gibco, 4.5g/L glucose), supplemented with 10% FCS and 1% PenStrep. Upon reaching ~70% confluency, medium was carefully aspirated from the wells and plates immediately placed on ice, and cells were resuspended in 1ml 75 mM ammonium carbonate pH 7.4, frozen in liquid nitrogen and stored at -80°C until metabolite extraction.

*Human haploid HAP1 cells and different culture medium compositions.* Cells (Horizon Discovery Group, United Kingdom) were cultured in 6-well plates in IMDM medium (Gibco, Thermo Fisher Scientific, USA), supplemented with 10% FCS and 1% PenStrep. Cells were passaged 1:10 every 2-3 days. Upon 70-80 % confluence, cells were rinsed twice with 75 mM ammonium carbonate (pH=7.4) and plates were immediately frozen in liquid nitrogen and stored at –80°C until metabolite extraction. To evaluate the influence of glucose and non-essential amino acids in the culture medium on the nucleotide sugar profile, IMDM culture medium was replaced with DMEM in various conditions, and cells were cultured for 8 hours before harvesting the cells for metabolite extraction. Glucose concentrations used were high (4.5 g/L), low (1 g/L), and 2.75 g/L. Cells were also cultured with and without 1% MEM Non-Essential Amino Acids (Gibco).

*C2C12 mouse myoblasts and differentiation to myotubes.* C2C12 myoblasts were cultured in 6-well plates on DMEM (Gibco, 4.5g/L glucose) supplemented with 10% FCS and 1% PenStrep. Upon reaching ~70% confluence, medium was refreshed and after 8 hours, the cells were washed and frozen as described for human fibroblasts. For differentiation to myotubes, cells were grown in 6-well plates until reaching 100% confluency. Differentiation was initiated by switching the growth medium to DMEM with low concentration of FCS (2%). During differentiation, medium was refreshed every day during 7 days. At 7 days, myotubes were washed twice quickly with 2 ml 75 mM ammonium carbonate pH 7.4 and snap-frozen using liquid nitrogen and stored at –80°C until metabolite extraction.

*Mouse B16-F10 melanoma cells and dynamic tracing of sialic acid synthesis in the presence of $3F_{ax}$-NeuNAc.* Mouse B16-F10 melanoma cells (ATCC CRL-6475) were cultured in MEM (Gibco) supplemented with 5% FBS (Greiner Bio-one), 1% MEM non-essential amino acids (Gibco), 0.15 % sodium bicarbonate (Gibco), 1 mM sodium pyruvate (Gibco), 1.5% MEM vitamins (Gibco) and 1x penicillin-streptomycin solution (Gibco). The cells were incubated in triplo for different time points (5 min, 1, 2, 4, 8, 24, and 48 hours) with medium containing PBS, 100 µM $Ac_5$NeuNAc or 100 µM $Ac_53F_{ax}$-NeuNAc, both synthesized as described before.[39] To correct for the addition of 1 mM [UL-$^{13}$C6]-GlcNAc in the subsequent experiment, 1 mM $^{12}$C-GlcNAc was added to each of these three conditions. For dynamic tracing of sialic acid biosynthesis in the presence of 100 µM $Ac_53F_{ax}$-NeuNAc, cells were incubated in triplo with 1 mM [UL-$^{13}$C6]-GlcNAc in the culture medium in combination with 100 µM $Ac_53F_{ax}$-NeuNAc or using 100 µM $Ac_5$NeuNAc or PBS as control, as detailed in Figure 4.

***Metabolite extraction -*** All samples were prepared in triplicate and for individual experiments extracted on the same day with identical timings and solvents. Frozen cell pellets as described above were extracted at -20˚C with cold 700 µl 2:2:1 (v/v/v) methanol:acetonitrile:water for 2 min. The supernatant was transferred to a separate vial and the extraction repeated with 700 µl cold extraction solvent for 3 min. The two extracts were pooled and centrifuged at 13,000 rpm for 3 minutes, using a pre-cooled centrifuge (4ºC). The resulting supernatants were dried using a vacuum centrifuge (SpeedVac, Thermo Fisher Scientific) at room temperature. The samples were reconstituted in 100 µl MilliQ and centrifuged for 3 min at 14,000 rpm. The supernatant was transferred to polypropylene (PP) autosampler vials or 96-well plates for LC-MS analysis, or stored at -80 °C until use.

***Organisms***

*Streptococcus Pneumonia.* The gram positive bacterium *Streptococcus pneumoniae* serotype 19A was grown on a blood agar plate (BD™ Columbia III Agar with 5% Sheep Blood 254098, containing 12g/L Pancreatic Digest of Casein, 5g/L Peptic Digest of Animal Tissue, 3g/L Yeast Extract, 3g/L Beef Extract, 1g/L Corn Starch, 5g/L Sodium Chloride, 13.5 g/L Agar 4g/L Growth factors, 5% Sheep Blood, Defibrinated, pH=7.3 ± 0.2), overnight, in a 37 °C, 5% $CO_2$ incubator. The next day, single colonies were inoculated into 30 ml THY broth (Difco 249240 and Labconsult SA-NV CON.1702; 3.1g/L Heart, Infusion from 500g, 20g/L Neopeptone, 2g/L Dextrose, 2g/L Sodium Chloride, 0.4g/L Disodium Phosphate, 2.5g/L Sodium Carbonate, 5g/L yeast) in a 50 ml tube, in a 37 °C, 5% $CO_2$ incubator, and grown until a optical density at a wavelength of 620 nm of 0.3. Then, samples were immediately placed onto ice with NaCl (Merck, 1064041000). In triplicate, 7 ml broth in a 15 ml tube was spun down by centrifugation (1 min, pre-cooled centrifuge at 4 °C, 3220 rcf). Supernatant was discarded and the pellet washed with 1.8 ml wash buffer (75 mM Ammonium Carbonate (Sigma 207861) in MilliQ, buffered with acetic acid (Sigma A6283) at pH 7.4 and cooled to 4 °C prior to use). The suspension was transferred to a 2 ml tube and spun down by centrifugation (1 minute, pre-cooled centrifuge at 4 °C, 25000 rcf). Supernatant was discarded and the pellet was stored at -80 °C until metabolite extraction. Metabolites were extracted at -20˚C with cold 1 ml 2:2:1 (v/v/v) methanol:acetonitrile:water for 5 min. Then this was centrifuged at 25000 rcf for 3 minutes, using a pre-cooled centrifuge at 4°C. The resulting supernatants were dried using a vacuum centrifuge at room temperature and the tubes stored at -80 °C until analysis.

*Haemophilus influenzae.* Non-typeable *Haemophilus influenzae* strain 86-028NP was grown in 7 mL Brain-Heart Infusion (BHI) medium (BD biosciences) supplemented with 1 µg/mL hemin and 2 µg/mL ß-nicotinamide adenine dinucleotide (Merck) to an optical density at 620 nm (OD620) of 0.5 in a 50 mL tube. Bacteria were pelleted by centrifugation with 3,220g for 10 minutes at 4°C. Bacterial pellet was suspended into 2 mL wash buffer (75 mM ammonium carbonate, pH 7.4) and pelleted by centrifugation with 3,220g for 10 minutes at 4°C. For metabolite extraction, the pellet was suspended into 1 mL 40:40:20 acetonitrile:methanol:water, transferred to a 1.5 mL tube and incubated 5 minutes at -20°C. Bacteria were pelleted by centrifugation at 16,100g for 3 minutes at 4°C. The supernatant was transferred to a new 1.5 mL tube and was dried in a vacuum centrifuge and the tubes stored at -80 °C until analysis.

*Yeast.* Fully $^{13}$C-labelled yeast metabolite extract was a kind gift of Dr. Gerrit Hermann (ISOtopic solutions, Austria) and used for analyses without further purification.

*Zebrafish.* Zebrafish (*Danio rerio*) were housed in recirculating systems on a 14/10 day-night regime. Husbandry was essentially performed as described by Lawrence.[50] Zebrafish embryos (3 days post fertilization; pool of 150 fish; n=3) and juvenile fish (4 weeks post fertilization; 1 fish/sample; n=4) were euthanized with a lethal dose of tricaine methanesulfonate (MS222). This study was approved by the Animal Ethics Committee of the University of Maastricht (Dier Experimenten Commissie, University of Maastricht) and the Dutch National Central Authority for Scientific Procedures on Animals (Centrale Commissie Dierproeven) (AVD107002016545). Care and use of animals were in agreement with the national and local guidelines. Zebrafish were suspended in 500 µL lysis buffer containing 75 mM ammonium carbonate buffer pH 7.4, homogenized in a potter tube (10 strokes) and subsequently sonicated by the ultrasonic processor UP50H (Hielscher; 2 mm diameter tip, amplitude 175 µm, power density 480 W/cm2). Lysates were centrifuged at 11,500 *g* for 20 minutes at 4 °C. Following protein analysis with the BCA Protein Assay Kit, 400 µg of the supernatant were thoroughly mixed with four volumes of cold methanol:acetronitrile (1:1) and incubated on ice for 10 minutes. After centrifugation at 14,000 *g* for 3 minutes at 4 °C, the supernatant was transferred to a new 1.5 mL tube and was dried in a vacuum centrifuge at room temperature. Tubes were stored at -80 °C until analysis.

*Drosophila melanogaster.* Flies were reared at normal fly food at 28°C. After 15 days, adult flies were collected (1-5 days of age), separated on gender and snap-frozen immediately in liquid nitrogen. For each polar metabolite extract, 10 flies were homogenized in 75mM ammonium carbonate (pH=7.4) using glass pestles. Protein concentrations were determined using the Bradford protein assay and an equivalent of 200ug protein was diluted to a total volume of 200uL using 75mM ammonium carbonate (pH=7.4). Polar metabolite extractions were performed by adding four volumes of cold 1:1 methanol:acetonitrile. After 5 minutes of extraction, samples were centrifuged at 13,000 rpm for 3 minutes at 4°C. Supernatants were dried using a vacuum centrifuge at room temperture and the tubes stored at -80 °C until analysis.

**Liquid chromatography – mass spectrometry analysis of sugar metabolites**

*UHPLC analysis of nucleotide sugars.* Separation of nucleotide sugars was achieved by ion pair-reverse phase chromatography using a modification of previously reported methodology.[30] An Agilent 1290 Infinity UHPLC system was used to inject 10 µL of cellular extract samples onto a HSS T3 column (Waters, 2.1 x 150 mm i.d., 1.8 µm particle size) that was maintained at a column temperature of 25 °C. Chromatography was performed using a 350 µL/min flow rate and a gradient from 0–10% mobile phase B over a 35-min total run time. Mobile phase A1 consisted of 20 mM triethylamine-acetic acid in $H_2O$, and mobile phase B1 consisted of 50% ACN/$H_2O$ (v/v). The gradient method was as follows (time: % B): 10.0 min: 0% B; 15 min: 4% B; 25 min: 4% B; 26 min: 10% B; 27 min: 10% B; 28 min: 0% B.

*UHPLC analysis for broad intracellular metabolome profiling.* Separation of a range of polar metabolites was achieved by an ion pair-reverse phase method as previously reported.[30] An Agilent 1290 Infinity UHPLC system was used to inject 10 µL of metabolite extracts onto a HSS T3 column (Waters, 2.1 x 150 mm i.d., 1.8 µm particle size) that was maintained at a column temperature of 40 ˚C. A gradient of mobile phases A2 (10 mM tributylamine, 15 mM acetic acid, 5% (v/v) methanol) and B2 (2-propanol) was used to separate the metabolites as indicated Table S3. The gradient method was as follows (time: % B): 9.5 min: 0% B; 14.5 min: 20% B; 20 min: 45% B; 27 min: 99% B; 31 min: 99% B; 31.5 min: 0% B; 32.0 min: 0% B. The flow rate was 0.25 mL/min, except between 31.5 and 32.0 min, where it was set to 0.15 mL/min.

The two LC methods were operated on the same LC system, by use of separate columns for the individual methods. Analysis of samples using both methods was performed via a switching valve, with switching between methods after each series of samples.

*QQQ Mass spectrometry and MRM acquisition.* An Agilent 6490 QQQ LC/MS system with a high-flow iFunnel ionization source, controlled by Agilent's MassHunter Workstation software (version B.05), was used for all LC-MRM/MS analyses. All acquisition methods used the following parameters: 3500 V capillary voltage and a 2000 V nozzle voltage, a sheath gas flow of 12 L/min (nitrogen) at a temperature of 200 °C, a drying gas flow of 15 L/min at a temperature of 200 °C, nebulizer gas flow at 20 psi, an MS operating pressure of 5 × 10−5 Torr, and Q1 and Q3 set to unit resolution (0.7 FWHM) and 10 ms dwell time. Agilent Optimizer software was used to find the optimal collision energies and fragment ions for each compound. A default 380 V fragmentor voltage and 4 V cell accelerator potential were used for all MRM ion pairs. The MRM acquisition method was settled with two ion pairs per nucleotide sugar, one quantifier and one qualifier transition (Table S1). Reported responses were based on the quantifier, which was the highest signal producing transition that has been verified to be free of interference. For isotopically labeled nucleotide sugars or synthetically modified nucleotide sugars, the MRM transitions were adapted based on fragmentation knowledge of the non-modified nucleotide sugars (Table S1). In addition to the 14 commercially available nucleotide sugars (Table S2), CDP-ribitol was synthesized and analyzed as described[33], CMP-NeuNGc was analyzed based on the conditions for CMP-NeuNAc with calculated mass difference, and UDP-ManNAc was analyzed using the conditions of UDP-GlcNAc. For the broad intracellular metabolome profiling method, dynamic MRM windows were used as reported in Table S3.

*Data analysis.* All MRM data were processed using Agilent MassHunter Quantitative Analysis software (Agilent B.07.00) with the Agilent Integrator algorithm for peak integration set with default values. All integrated peaks were manually inspected to ensure correct peak detection and accurate integration. For data interpretation of nucleotide sugars, normalization was performed of peak area over the total peak area of all nucleotide sugars to obtain relative abundances. Differential analyses were performed between conditions and visualized using GraphPad Prism 5.03 or Microsoft Office Excel 2007.

For dynamic tracing of isotopically labeled *N*-acetylglucosamine, the abundance of labelled fraction of each investigated metabolite was calculated using the ratio of $^{13}C/^{12}C$ isotopes of a specific metabolite.

**Determination of linearity, limit of detection (LOD) and limit of quantitation (LOQ) for nucleotide sugars in fibroblasts**

For 14 nucleotide sugars (Table S2) that we detected in human cell lines and for which commercial standards were available, calibration curves were generated to determine the dynamic range by linear regression analysis. A mix of 10 control primary dermal fibroblast extracts was used for the dynamic range determination. Each sample was spiked with an increasing amount of the 14 nucleotide sugars, spanning a 10000-fold range. The linear range (Table S2) was based on the level of endogenous nucleotide sugars, with the lower standard points designed primarily to determine the LOQ. For determination of the LOD, a signal/noise ratio of 3 was taken. For determination of the LOQ, 10 times a blanc injection was performed. The LOQ was calculated by taking the signal of the standards from the blanc injection plus 10 times the standard deviation of the noise.

**Structural modelling of CMAS and GNE**

For structural modeling of CMAS, PDB file 1qwj (X-ray structure of murine CMAS[41]) was used to model the interaction with $3F_{ax}$-NeuNAc and NeuNAc in the active site of the CMAS enzyme. NeuNAc was redocked and $3F_{ax}$-NeuNAc was docked in the A site of the AB dimer.[41] The affinities of NeuNAc and $3F_{ax}$-NeuNAc were predicted to be 50.8 and 49.8 µM, respectively. For structural modeling of GNE, PDB file 4ZHT[42] was used to model the interaction of CMP-$3F_{ax}$-NeuNAc and CMP-NeuNAc with the allosteric binding site of human GNE. CMP-$3F_{ax}$-NeuNAc and CMP-NeuNAc bound to the allosteric binding site of GNE with calculated affinities of 2.4 µM and 0.6 µM, respectively. All modelling was performed with Yasara 17.12.24.[51] Local docking with ligand flexibility was performed 25 times with the default Yasara macro dock_runlocal.mcr modified to use the AutoDockLS algorithm.

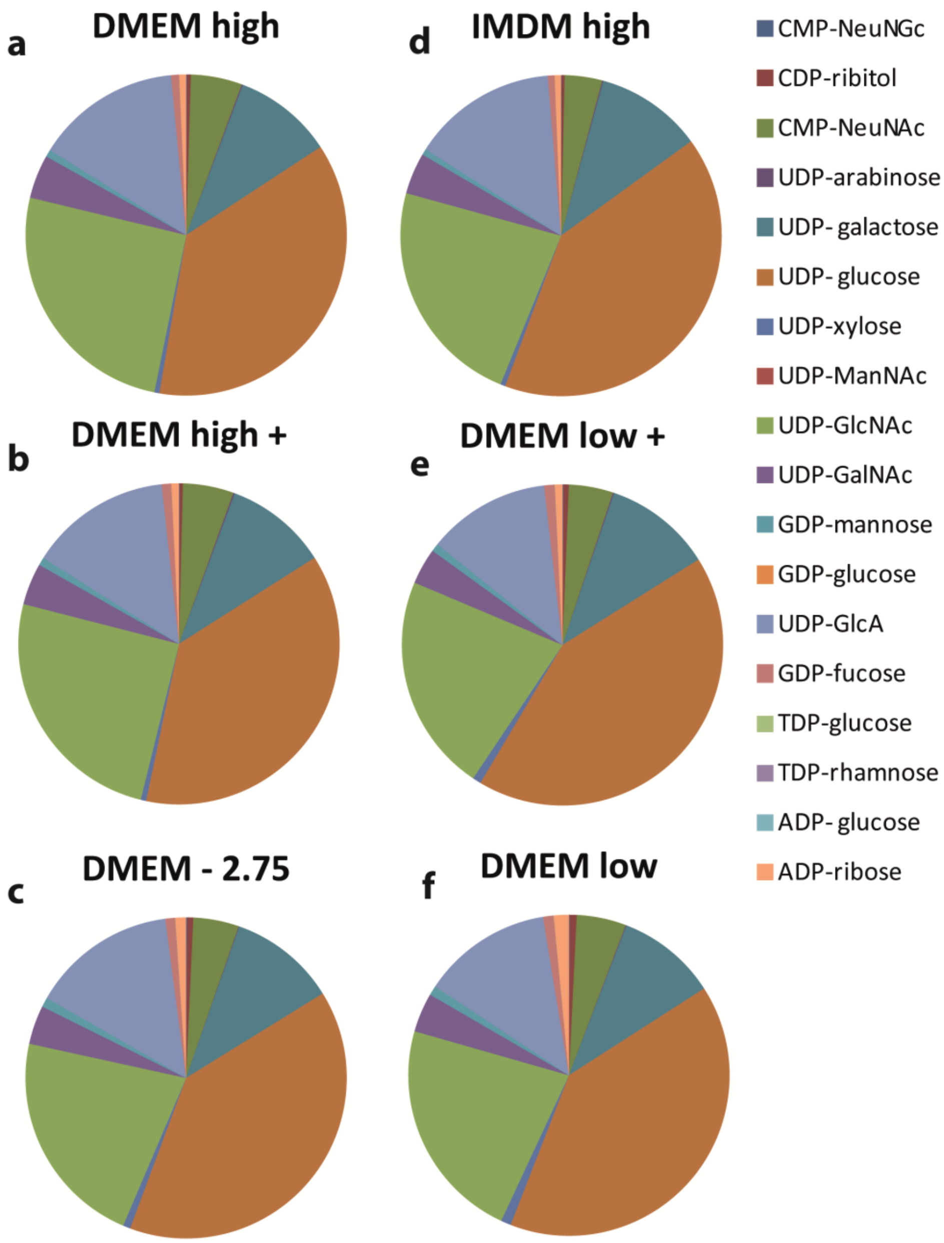


**Supplementary Figure 1. The effect of different media compositions on nucleotide sugar profiles in HAP1 cells.** Human haploid (HAP1) cells were grown in standard high-glucose IMDM medium, then switched to DMEM with different media compositions and cultured for 8 hours, after which polar metabolites were extracted. Different conditions included glucose concentration (4.5 g/L – high; 1 g/L – low; and 2.75 gr/L) and the presence ('+') or absence of 1% MEM Non-Essential Amino Acids. Experiments were performed in triplo.

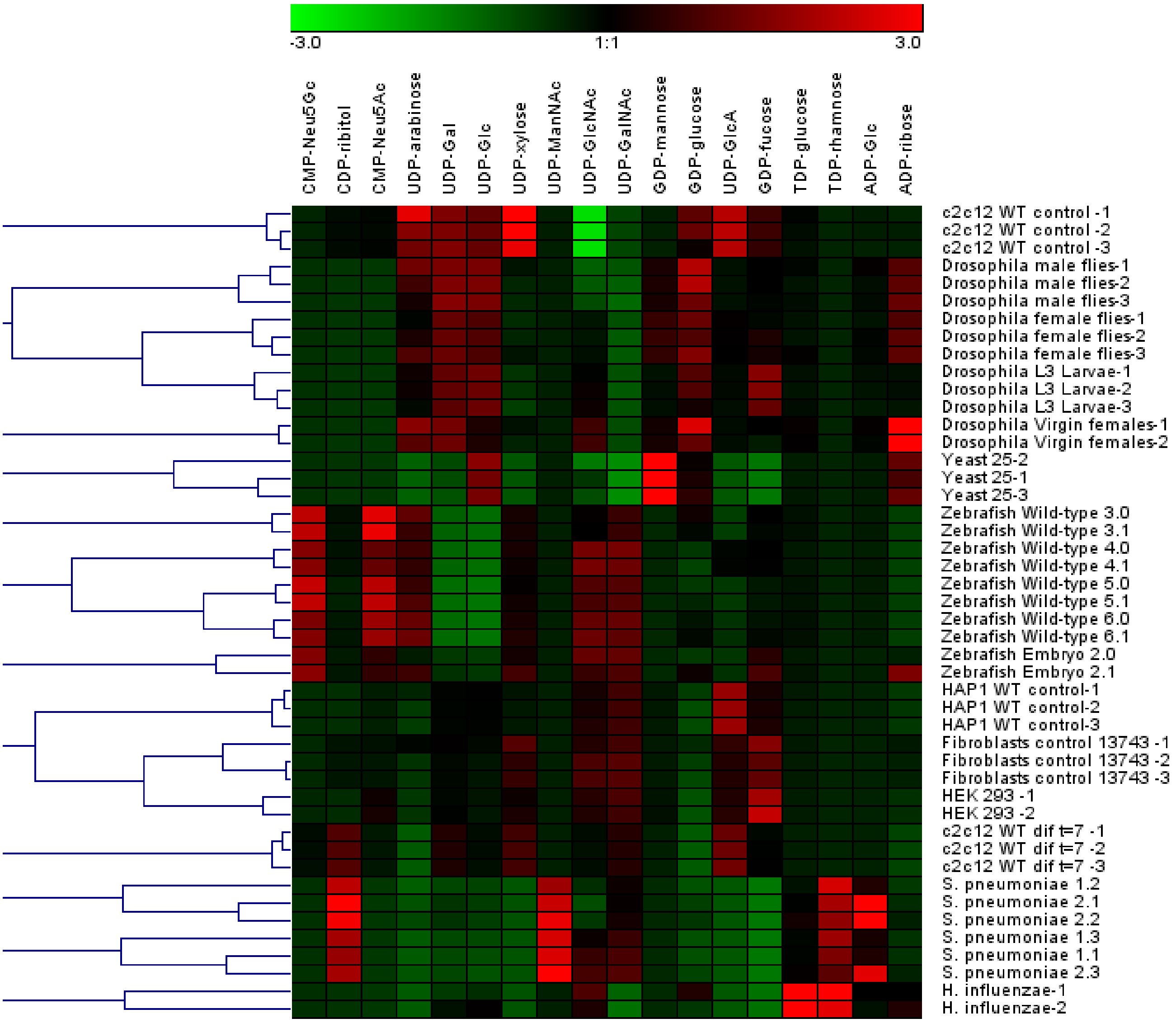


**Supplementary Figure 2. Hierarchical clustering of the 9 different model organisms and cell lines.** 2D hierarchical clustering (HCL) was performed on the MRM data of in total 46 samples of organisms and cell lines, consisting of the relative abundances of 18 nucleotide sugars. Genesis version 1.7.7 software was used[52] based on average linkage clustering and Pearson correlations.

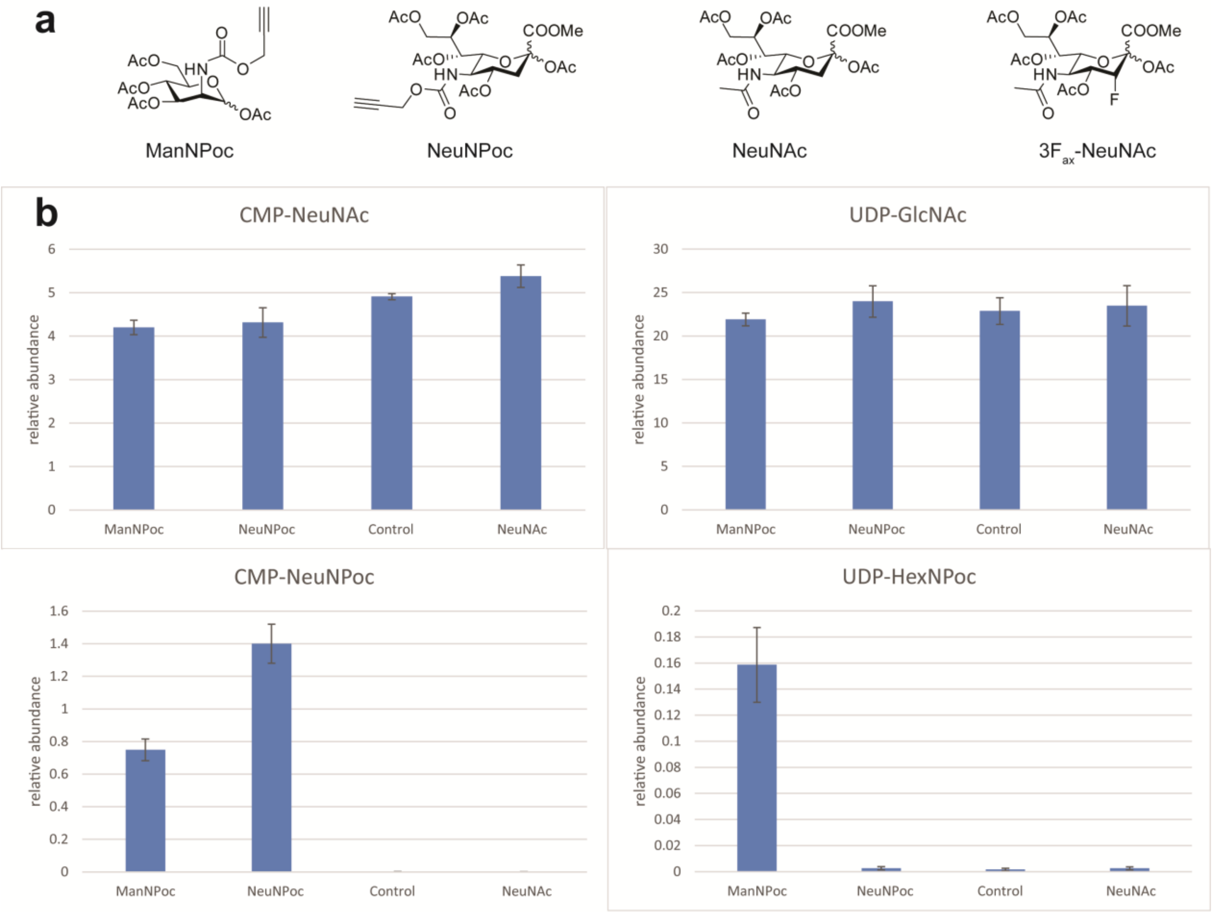


**Supplementary Figure 3. Metabolic insights into the uptake of Poc derivatives in fibroblasts.** a) Chemical structures of the ManNPoc and NeuNPoc derivatives, sialic acid and its fluorinated derivative as used in this study. b) Relative abundances of CMP-NeuNAc, UDP-GlcNAc, CMP-NeuNPoc and UDP-HexNPoc after 48 hours incubation with respectively 100 µM $Ac_5$ManNPoc, 100 µM $Ac_5$NeuNPoc or 100 µM $Ac_5$NeuNAc. Control fibroblasts incubated with PBS were taken as a control. The average and standard deviation are plotted of triplicate samples.

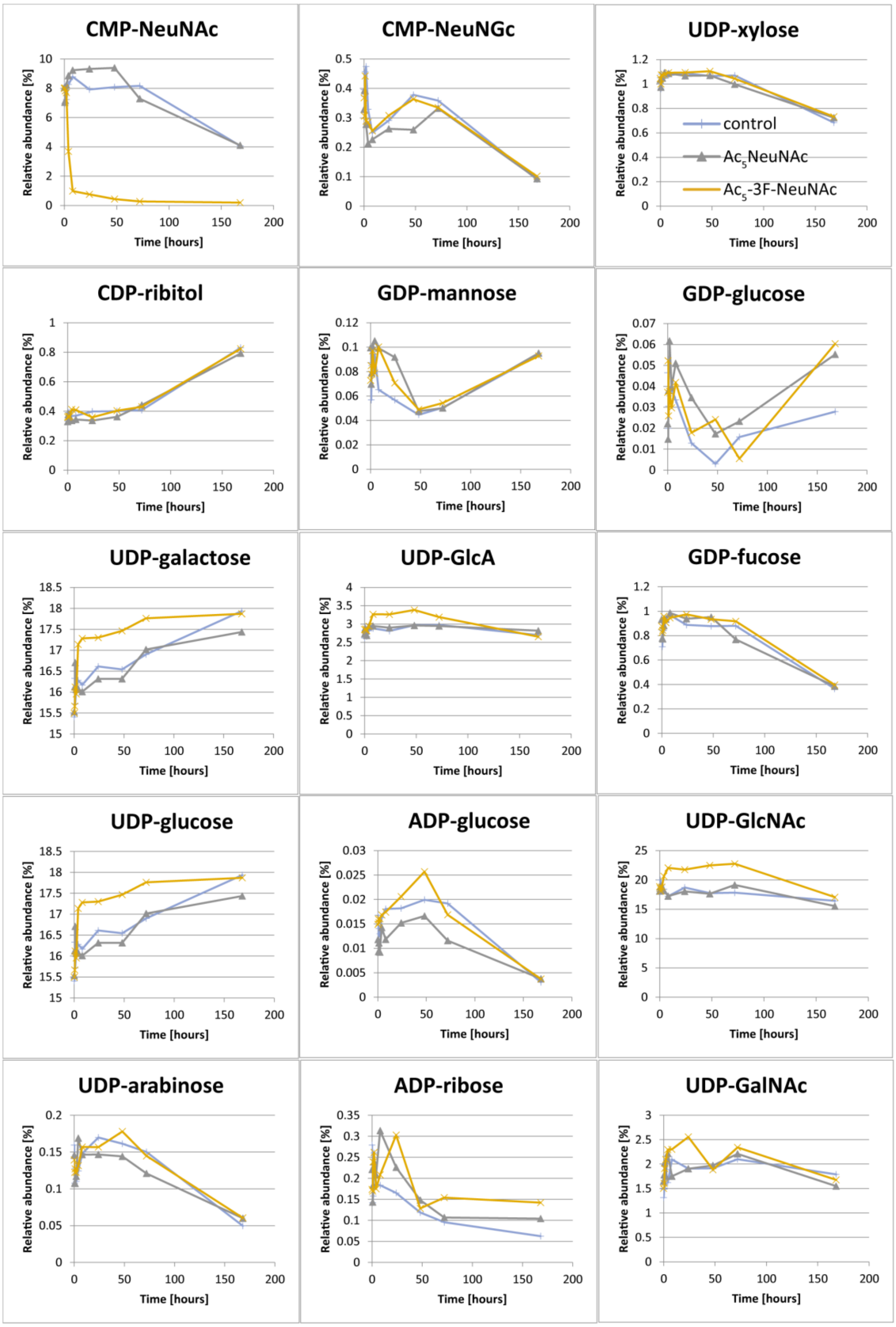


**Supplementary Figure 4. Overview of the effect of $3F_{ax}$-NeuNAc in B16-F10 cells on all nucleotide sugars in time.** The relative abundances of nucleotide sugars are shown in time after incubation of B16-F10 cells with $Ac_5 3F_{ax}$-NeuNAc (orange line), $Ac_5$NeuNAc (green line) or PBS (blue line). Experimental conditions are described in the methods section.
