## Supplemental Table 1 for "Dynamic analysis of sugar metabolism reveals the mechanisms of action of synthetic sugar analogs"

### Supplementary Table 1. MRM transitions and collision energies of nucleotide sugars and their isotopically labeled or synthetic analogs.

| Compound | Chemical formula | Retention time (r | dMRM time wind | monoisotopic mas | Q1 mass (amu) | Q3 mass (amu) | collision energy (eV) |
|---|---|---|---|---|---|---|---|
| UDP-xylose | $C_{14}H_{22}N_2O_{16}P_2$ | 9,6 | 1 | 536,04 | 535,03 | 79.1 | 30 |
| UDP-xylose | | | 1 | | | 323.2 | 20 |
| 13C5-UDP-xylose | 13C5-$C_9H_{22}N_2O_{16}P_2$ | | 1 | 541,04 | 540,03 | 79.1 | 30 |
| 13C5-UDP-xylose | | | 1 | | | 323.2 | 20 |
| CDP-ribitol | $C_{14}H_{25}N_3O_{15}P_2$ | 4,8 | 1 | 537,08 | 536,07 | 78.8 | 50 |
| CDP-ribitol | | | 1 | | | 322 | 20 |
| 13C5-CDP-ribitol | 13C5-$C_9H_{25}N_3O_{15}P_2$ | | 1 | 542,08 | 541,07 | 78.8 | 50 |
| 13C5-CDP-ribitol | | | 1 | | | 322 | 20 |
| UDP-galactose | $C_{15}H_{24}N_2O_{17}P_2$ | 7,4 | 1 | 566,06 | 565,05 | 79 | 30 |
| UDP-galactose | | | 1 | | | 323.1 | 24 |
| 13C6-UDP-galactose | 13C6-$C_9H_{24}N_2O_{17}P_2$ | | 1 | 572,06 | 571,05 | 79 | 30 |
| 13C6-UDP-galactose | | | 1 | | | 323.1 | 24 |
| UDP-glucose | $C_{15}H_{24}N_2O_{17}P_2$ | 8,3 | 1 | 566,06 | 565,05 | 79 | 44 |
| UDP-glucose | | | 1 | | | 323.2 | 20 |
| 13C6-UDP-gluose | 13C6-$C_9H_{24}N_2O_{17}P_2$ | | 1 | 572,06 | 571,05 | 79 | 44 |
| 13C6-UDP-glucose | | | 1 | | | 323.2 | 20 |
| UDP-GalNAc | $C_{17}H_{27}N_3O_{17}P_2$ | 10,5 | 1 | 607,08 | 606,07 | 78.9 | 50 |
| UDP-GalNAc | | | 1 | | | 273 | 36 |
| 13C6-UDP-GalNAc | 13C6-$C_{11}H_{27}N_3O_{17}P_2$ | | 1 | 613,08 | 612,07 | 78.9 | 50 |
| 13C6-UDP-GalNAc | | | 1 | | | 273 | 36 |
| UDP-GlcNAc | $C_{17}H_{27}N_3O_{17}P_2$ | 10,7 | 1 | 607,08 | 606,07 | 282.1 | 30 |
| UDP-GlcNAc | | | 1 | | | 385.1 | 28 |
| 13C6-UDP-GlcNAc | 13C6-$C_{11}H_{27}N_3O_{17}P_2$ | | 1 | 613,08 | 612,07 | 282.1 | 30 |
| 13C6-UDP-GlcNAc | | | 1 | | | 385.1 | 28 |
| UDP-HexNPoc | $C_{19}H_{27}N_3O_{18}P_2$ | 18,0 | 1 | 647,08 | 646,07 | 282.1 | 30 |
| UDP-HexNPoc | | | | | | | |
| CMP-NeuNAc | $C_{20}H_{31}N_4O_{16}P$ | 6,3 | 1 | 614,15 | 613,14 | 78.9 | 54 |
| CMP-NeuNAc | | | 1 | | | 321.9 | 20 |
| 13C6-CMP-NeuNAc | 13C6-$C_{14}H_{31}N_4O_{16}P$ | | 1 | 620,15 | 619,14 | 78.9 | 54 |
| 13C6-CMP-NeuNAc | | | 1 | | | 321.9 | 20 |
| CMP-3Fax-NeuNAc | $C_{20}H_{30}FN_4O_{16}P$ | 5,8 | 1 | 632,13 | 631,13 | 78.9 | 54 |
| CMP-3Fax-NeuNAc | | | 1 | | | 321.9 | 20 |
| CMP-NeuNPoc | $C_{22}H_{31}N_4O_{17}P$ | 16,2 | 1 | 654,13 | 653,13 | 78.9 | 54 |
| CMP-NeuNPoc | | | 1 | | | 321.9 | 20 |
| CMP-NeuNGc | $C_{20}H_{31}N_4O_{17}P$ | | 1 | 630,14 | 629,13 | 78.9 | 54 |
| CMP-NeuNGc | | | 1 | | | 321.9 | 20 |
| 13C6-CMP-NeuNGc | 13C6-$C_{14}H_{31}N_4O_{17}P$ | | 1 | 636,14 | 635,13 | 78.9 | 54 |
| 13C6-CMP-NeuNGc | | | 1 | | | 321.9 | 20 |
| GDP-glucose | $C_{16}H_{25}N_5O_{16}P_2$ | 14,9 | 2 | 605,08 | 604,07 | 362 | 24 |
| GDP-glucose | | | 2 | | | 78.9 | 44 |
| 13C6-GDP-glucose | 13C6-$C_{10}H_{25}N_5O_{16}P_2$ | | 2 | 611,08 | 610,07 | 362 | 24 |
| 13C6-GDP-glucose | | | 2 | | | 78.9 | 44 |
| GDP-mannose | $C_{16}H_{25}N_5O_{16}P_2$ | 13,6 | 2 | 605,08 | 604,07 | 424 | 32 |
| GDP-mannose | | | 2 | | | 78.8 | 68 |
| 13C6-GDP-mannose | 13C6-$C_{10}H_{25}N_5O_{16}P_2$ | | 2 | 611,08 | 610,07 | 78.8 | 68 |
| 13C6-GDP-mannose | | | 2 | | | 424 | 32 |
| GDP-fucose | $C_{16}H_{25}N_5O_{15}P_2$ | 17,1 | 2 | 589,08 | 588,07 | 442 | 20 |
| GDP-fucose | | | 2 | | | 424 | 28 |
| GDP-fucose | | | 2 | | | 78.9 | 50 |
| 13C6-GDP-fucose | 13C6-$C_{10}H_{25}N_5O_{15}P_2$ | | 2 | 595,08 | 594,07 | 442 | 20 |
| 13C6-GDP-fucose | | | 2 | | | 424 | 28 |
| 13C6-GDP-fucose | | | 2 | | | 78.9 | 50 |
| ADP-glucose | $C_{16}H_{25}N_5O_{15}P_2$ | 19,9 | 2 | 589,08 | 588,07 | 346.2 | 24 |
| ADP-glucose | | | 2 | | | 241.1 | 28 |
| 13C6-ADP-glucose | 13C6-$C_{10}H_{25}N_5O_{15}P_2$ | | 2 | 595,08 | 594,07 | 346.2 | 24 |
| 13C6-ADP-glucose | | | 2 | | | 241.1 | 28 |
| UDP-GlcA | $C_{15}H_{22}N_2O_{18}P_2$ | 15,4 | 2 | 580,03 | 579,02 | 403 | 20 |
| UDP-GlcA | | | 2 | | | 158.9 | 42 |
| 13C6-UDP-GlcA | 13C6-$C_9H_{22}N_2O_{18}P_2$ | | 2 | 586,03 | 585,02 | 403 | 20 |
| 13C6-UDP-GlcA | | | 2 | | | 158.9 | 42 |
| ADP-ribose | $C_{15}H_{23}N_5O_{14}P_2$ | 20,0 | 2 | 559,07 | 558,06 | 346.2 | 24 |
| ADP-ribose | | | 2 | | | 78.8 | 64 |
| 13C5-ADP-ribose | 13C5-$C_{10}H_{23}N_5O_{14}P_2$ | | 2 | 564,07 | 563,06 | 346.2 | 24 |
| 13C5-ADP-ribose | | | 2 | | | 78.8 | 64 |
| TDP-glucose | $C_{16}H_{26}N_2O_{16}P_2$ | 18,8 | 2 | 564,08 | 563,07 | 321 | 20 |
| TDP-glucose | | | 2 | | | 79.1 | 48 |
| 13C6-TDP-glucose | 13C6-$C_{10}H_{26}N_2O_{16}P_2$ | | 2 | 570,08 | 569,07 | 321 | 20 |
| 13C6-TDP-glucose | | | 2 | | | 79.1 | 48 |
| TDP-rhamnose | $C_{16}H_{26}N_2O_{15}P_2$ | 19,3 | 2 | 548,08 | 547,07 | 321 | 20 |
| TDP-rhamnose | | | 2 | | | 79.1 | 48 |
| 13C6-TDP-rhamnose | 13C6-$C_{10}H_{26}N_2O_{15}P_2$ | | 2 | 554,08 | 553,07 | 321 | 20 |
| 13C6-TDP-rhamnose | | | 2 | | | 79.1 | 48 |
| UDP-Arabinose | $C_{14}H_{22}N_2O_{16}P_2$ | 7,1 | 1 | 536,04 | 535,03 | 322,9 | 20 |
| UDP-Arabinose | | | 1 | | | 78,9 | 30 |
| 13C5-UDP-Arabinose | 13C5-$C_9H_{22}N_2O_{16}P_2$ | | 1 | 541,04 | 540,03 | 322,9 | 20 |
| 13C5-UDP-Arabinose | | | 1 | | | 78,9 | 30 |
| UDP-ManNAc | $C_{17}H_{27}N_3O_{17}P_2$ | 18,1 | 1 | 607,08 | 606,07 | 78.9 | 50 |
| | | | | | | 273 | 36 |
