## Supplemental Table 2 for "Dynamic analysis of sugar metabolism reveals the mechanisms of action of synthetic sugar analogs"

**Supplementary Table 2. Acquisition parameters and results of validation for 14 nucleotide sugars.**

| Compound | HMDB ID | retention time (min) | lower limit of detection (µM) | lower limit of quantification (µM) | linear range (log3) | monoisotopic mass (amu) | Q1 mass (amu) | Q3 mass (amu) | collision energy (eV) | Kolom1 |
|---|---|---|---|---|---|---|---|---|---|---|
| CMP-NeuNAc | 01176 | 6,3 | 0,001 | 0,003 | 3 nM - 3,13 µM | 614,15 | 613,1 | 321,9 | 20 | Quan |
| | | | | | | | | 78,9 | 54 | Qual |
| UDP-Galactose | 00302 | 7,4 | 0,001 | 0,003 | 3 nM - 6,25 µM | 566,06 | 565,1 | 323,1 | 24 | Quan |
| | | | | | | | | 79 | 30 | Qual |
| UDP-Glucose | 00286 | 8,3 | 0,001 | 0,003 | 3 nM - 6,25 µM | 566,06 | 565,1 | 323,2 | 20 | Quan |
| | | | | | | | | 79 | 44 | Qual |
| UDP-GalNAc | 00304 | 10,5 | 0,009 | 0,03 | 31 nM - 3,13 µM | 607,08 | 606,1 | 273 | 36 | Quan |
| | | | | | | | | 78,9 | 50 | Qual |
| UDP-GlcNAc | 00290 | 10,7 | 0,003 | 0,009 | 9 nM - 3,13 µM | 607,08 | 606,1 | 385,1 | 28 | Quan |
| | | | | | | | | 282,1 | 30 | Qual |
| GDP-mannose | 01163 | 13,6 | 0,003 | 0,009 | 9 nM - 3,13 µM | 605,08 | 604,1 | 424 | 32 | Quan |
| | | | | | | | | 78,8 | 68 | Qual |
| TDP-glucose | 01328 | 18,8 | 0,003 | 0,009 | 9 nM - 1,25 µM | 564,08 | 563,1 | 321 | 20 | Quan |
| | | | | | | | | 79,1 | 48 | Qual |
| GDP-glucose | 03351 | 14,9 | 0,003 | 0,009 | 9 nM - 3,13 µM | 605,08 | 604,1 | 362 | 24 | Quan |
| | | | | | | | | 78,9 | 44 | Qual |
| GDP-fucose | 01095 | 17,1 | 0,003 | 0,009 | 9 nM - 1,25 µM | 589,08 | 588,1 | 442 | 20 | Quan |
| | | | | | | | | 424 | 28 | Qual |
| | | | | | | | | 78,9 | 50 | extra |
| ADP-glucose | 06557 | 19,9 | 0,003 | 0,009 | 9 nM - 9,375 µM | 589,08 | 588,1 | 346,2 | 24 | Quan |
| | | | | | | | | 241,1 | 28 | Qual |
| ADP-ribose | 01178 | 20 | 0,006 | 0,02 | 22 nM - 12,5 µM | 559,07 | 558,1 | 346,2 | 24 | Quan |
| | | | | | | | | 78,8 | 64 | Qual |
| UDP-xylose | 01018 | 9,6 | 0,003 | 0,009 | 9 nM - 3,13 µM | 536,04 | 535 | 323,2 | 20 | Quan |
| | | | | | | | | 79,1 | 30 | Qual |
| UDP-arabinose | 12303 | 7,1 | 0,001 | 0,003 | 3 nM - 3,13 µM | 536,04 | 535 | 322,9 | 20 | Quan |
| | | | | | | | | 78,9 | 30 | Qual |
| UDP-GlcA | 00935 | 15,4 | 0,001 | 0,003 | 3 nM - 9,375 µM | 580,03 | 579 | 403 | 20 | Quan |
| | | | | | | | | 158,9 | 42 | Qual |
