## Supplemental Table 3 for "Dynamic analysis of sugar metabolism reveals the mechanisms of action of synthetic sugar analogs"

**Supplementary Table S3. Acquisition parameters of metabolites as analyzed in the TBA method.**

| Compound name | Chemical formula | HMDB ID | retention time (min) | monoisotopic mass (amu) | Q1 mass (amu) | Q3 mass (amu) | collision energy (eV) |
|---|---|---|---|---|---|---|---|
| Acetyl coenzyme A | C23H38N7O17P3S | HMDB0001206 | 24.0 | 809,13 | 808.10 | 408.00 | 36 |
| N-Acetyl-D-galactosamine | C8H15NO6 | HMDB0000212 | 1.5 | 221,09 | 220.1 | 119.00 | 0 |
| N-Acetyl-D-galactosamine 6-P | C8H16NO9P | HMDB0059626 | 14,2 | 301,06 | 300.00 | 97.00 | 16 |
| N-Acetyl-D-galactosamine 6-P | C8H16NO9P | HMDB0059626 | 14,2 | 301,06 | 300.00 | 79.00 | 36 |
| N-Acetyl-D-glucosamine | C8H15NO6 | HMDB0000215 | 1.5 | 221,09 | 220.1 | 59.00 | 20 |
| N-Acetyl-D-glucosamine 1-P | C8H16NO9P | HMDB0001367 | 14.0 | 301,06 | 300.00 | 97.00 | 18 |
| N-Acetyl-D-glucosamine 1-P | C8H16NO9P | HMDB0001367 | 14.0 | 301,06 | 300.00 | 79.00 | 48 |
| N-Acetyl-D-glucosamine 6-P | C8H16NO9P | HMDB0001062 | 13.8 | 301,06 | 300.00 | 97.00 | 16 |
| N-Acetyl-D-glucosamine 6-P | C8H16NO9P | HMDB0001062 | 13.8 | 301,06 | 300.00 | 79.00 | 36 |
| N-Acetyl-D-mannosamine | C8H15NO6 | HMDB0001129 | 1.5 | 221,09 | 220.1 | 59.00 | 0 |
| N-Acetyl-D-mannosamine 6-P | C8H16NO9P | HMDB0062500 | 14.5 | 301,06 | 300.1 | 87.10 | 9 |
| N-Acetyl-D-mannosamine 6-P | C8H16NO9P | HMDB0062500 | 14.5 | 301,06 | 300.1 | 78.90 | 41 |
| N-Acetylneuraminic acid | C11H19NO9 | HMDB0000230 | 7.0 | 309,11 | 308.1 | 169.90 | 10 |
| N-Acetylneuraminic acid | C11H19NO9 | HMDB0000230 | 7.0 | 309,11 | 308.1 | 87.00 | 14 |
| N-Acetylneuraminic acid 9-P | C13H21NO10 | HMDB0000794 | 20.9 | 351,12 | 388.1 | 79.00 | 41 |
| Adenine | C5H5N5 | HMDB0000034 | 2.9 | 135,05 | 134.00 | 107.00 | 18 |
| Adenine | C5H5N5 | HMDB0000034 | 2.9 | 135,05 | 134.00 | 92.1 | 20 |
| Adenosine 5-monophosphate | C10H14N5O7P | HMDB0000045 | 19.0 | 347,06 | 346.00 | 97.00 | 24 |
| Adenosine 5-monophosphate | C10H14N5O7P | HMDB0000045 | 19.0 | 347,06 | 346.00 | 79.00 | 38 |
| Adenosine 5-diphosphate | C10H15N5O10P2 | HMDB0001341 | 22.0 | 427,03 | 426.0 | 328.0 | 16 |
| Adenosine 5-diphosphate | C10H15N5O10P2 | HMDB0001341 | 22.0 | 427,03 | 426.0 | 159.0 | 28 |
| Adenosine 5-triphosphate | C10H16N5O13P3 | HMDB0000538 | 23.2 | 507,00 | 506.00 | 408.10 | 22 |
| Adenosine 5-triphosphate | C10H16N5O13P3 | HMDB0000538 | 23.2 | 507,00 | 506.00 | 159.00 | 38 |
| L-Arabinose | C5H10O5 | HMDB0000646 | 3.0 | 150,05 | 149.00 | 89.10 | 4 |
| L-Arabinose | C5H10O5 | HMDB0000646 | 3.0 | 150,05 | 149.00 | 59.20 | 12 |
| L-Arabitol | C5H12O5 | HMDB0001851 | 1.5 | 152,07 | 151.00 | 89.00 | 12 |
| L-Arabitol | C5H12O5 | HMDB0001851 | 1.5 | 152,07 | 151.00 | 71.02 | 16 |
| CDP-ribitol | C14H25N3O15P2 | n.a. | 20.5 | 537,08 | 536.1 | 322.0 | 20 |
| CDP-ribitol | C14H25N3O15P2 | n.a. | 20.5 | 537,08 | 536.1 | 78.8 | 50 |
| CMP-NeuNAc | C20H31N4O16P | HMDB0001176 | 20.5 | 614,15 | 613.14 | 321.90 | 20 |
| CMP-NeuNAc | C20H31N4O16P | HMDB0001176 | 20.5 | 614,15 | 613.14 | 78.90 | 54 |
| Cytidine | C9H13N3O5 | HMDB0000089 | 1.8 | 243,09 | 242.10 | 109.00 | 8 |
| Cytidine | C9H13N3O5 | HMDB0000089 | 1.8 | 243,09 | 242.10 | 42.20 | 16 |
| Cytidine 5-monophosphate | C9H14N3O8P | HMDB0000095 | 15.5 | 323,05 | 322.00 | 97.00 | 22 |
| Cytidine 5-monophosphate | C9H14N3O8P | HMDB0000095 | 15.5 | 323,05 | 322.00 | 79.00 | 44 |
| Cytidine 5-diphosphate | C9H15N3O11P2 | HMDB0001546 | 21.0 | 403,02 | 402.01 | 158.92 | 24 |
| Cytidine 5-diphosphate | C9H15N3O11P2 | HMDB0001546 | 21.0 | 403,02 | 402.01 | 79.00 | 48 |
| Cytidine 5-triphosphate | C9H16N3O14P3 | HMDB0000082 | 23.0 | 482,98 | 482.00 | 158.80 | 40 |
| Cytidine 5-triphosphate | C9H16N3O14P3 | HMDB0000082 | 23.0 | 482,98 | 482.00 | 79.00 | 40 |
| D-Erythrose 4-P | C4H9O7P | HMDB0001321 | 14.0 | 200,01 | 199.00 | 97.00 | 14 |
| D-Erythrose 4-P | C4H9O7P | HMDB0001321 | 14.0 | 200,01 | 199.00 | 79.00 | 48 |
| D-Fructose 1-P | C6H12O6 | HMDB0062538 | 14.0 | 180,06 | 259.00 | 97.00 | 14 |
| D-Fructose 1-P | C6H12O6 | HMDB0062538 | 14.0 | 180,06 | 259.00 | 79.00 | 48 |
| D-Fructose 6-P | C6H13O9P | HMDB0000124 | 13.5 | 260,03 | 259.00 | 97.00 | 14 |
| D-Fructose 6-P | C6H13O9P | HMDB0000124 | 13.5 | 260,03 | 259.00 | 79.00 | 48 |
| D-Fructose 1,6-bisphosphate | C6H14O12P2 | HMDB0001058 | 20.3 | 340,00 | 338.9 | 241.01 | 12 |
| D-Fructose 1,6-bisphosphate | C6H14O12P2 | HMDB0001058 | 20.3 | 340,00 | 338.9 | 97.00 | 22 |
| Galactitol | C6H14O6 | HMDB0000107 | 1.4 | 182,08 | 181.07 | 101.00 | 13 |
| Galactitol | C6H14O6 | HMDB0000107 | 1.4 | 182,08 | 181.07 | 59.00 | 21 |
| Galactonic acid | C6H12O7 | HMDB0000565 | 4.0 | 196,06 | 195.10 | 129.00 | 11 |
| Galactonic acid | C6H12O7 | HMDB0000565 | 4.0 | 196,06 | 195.10 | 75.10 | 18 |
| D-Galactose 1-P | C6H13O9P | HMDB0000645 | 13.4 | 260,03 | 259.00 | 240.90 | 9 |
| D-Galactose 1-P | C6H13O9P | HMDB0000645 | 13.4 | 260,03 | 259.00 | 79.00 | 28 |
| Glucose | C6H12O6 | HMDB0000122 | 1.4 | 180,06 | 179.10 | 89.10 | 5 |
| Glucose | C6H12O6 | HMDB0000122 | 1.4 | 180,06 | 179.10 | 59.00 | 17 |
| D-Glucose 1-P | C6H13O9P | HMDB0001586 | 13.6 | 260,03 | 259.00 | 240.90 | 9 |
| D-Glucose 1-P | C6H13O9P | HMDB0001586 | 13.6 | 260,03 | 259.00 | 79.00 | 28 |
| D-Glucose 6-P | C6H13O9P | HMDB0001401 | 12.9 | 260,03 | 259.00 | 97.00 | 14 |
| D-Glucose 6-P | C6H13O9P | HMDB0001401 | 12.9 | 260,03 | 259.00 | 79.00 | 48 |
| Glucuronic acid | C6H10O7 | HMDB0000127 | 5.0 | 194,04 | 193.00 | 113.00 | 14 |
| L-Glutamine | C5H10N2O3 | HMDB0000641 | 1.4 | 146,07 | 145.10 | 127.00 | 7 |
| L-Glutamine | C5H10N2O3 | HMDB0000641 | 1.4 | 146,07 | 145.1 | 109.00 | 10 |
| Glyceraldehyde 3-P | C3H7O6P | HMDB0001112 | 14.0 | 170,00 | 169.00 | 97.00 | 4 |
| Glyceraldehyde 3-P | C3H7O6P | HMDB0001112 | 14.0 | 170,00 | 169.00 | 79.00 | 28 |
| N-Glycolylneuraminic acid | C11H19NO10 | HMDB0000833 | 6.0 | 325,10 | 324.09 | 116.00 | 13 |
| N-Glycolylneuraminic acid | C11H19NO10 | HMDB0000833 | 6.0 | 325,10 | 324.09 | 87.10 | 25 |
| Guanine | C5H5N5O | HMDB0000132 | 3.0 | 151,05 | 150.00 | 133.02 | 12 |
| Guanine | C5H5N5O | HMDB0000132 | 3.0 | 151,05 | 150.00 | 66.01 | 36 |
| Guanosine | C10H13N5O5 | HMDB0000133 | 8.8 | 283,09 | 282.10 | 150.00 | 17 |
| Guanosine | C10H13N5O5 | HMDB0000133 | 8.8 | 283,09 | 282.10 | 132.90 | 32 |
| Mannitol | C6H14O6 | HMDB0000765 | 14.0 | 182,08 | 181.1 | 71.00 | 22 |
| D-Mannose | C6H12O6 | HMDB0000169 | 1.4 | 180,06 | 179.10 | 89.00 | 4 |
| D-Mannose | C6H12O6 | HMDB0000169 | 1.4 | 180,06 | 179.10 | 59.20 | 16 |
| D-Mannose 1-P | C6H13O9P | HMDB0006330 | 13.5 | 260,03 | 259.00 | 79.00 | 28 |
| D-Mannose 6-P | C6H13O9P | HMDB0001078 | 13.1 | 260,03 | 259.00 | 97.00 | 14 |
| D-Mannose 6-P | C6H13O9P | HMDB0001078 | 13.1 | 260,03 | 259.00 | 79.00 | 48 |
| Phosphoenolpyruvic acid | C3H5O6P | HMDB0000263 | 21.7 | 167,98 | 167.00 | 79.00 | 12 |
| 6-Phosphogluconic acid | C6H13O10P | HMDB0001316 | 14.0 | 276,02 | 275.0 | 79.0 | 50 |
| 5-Phosphoribosyl 1-PP | C5H13O14P3 | HMDB0000280 | 16.0 | 389,95 | 388.90 | 291.00 | 9 |
| 5-Phosphoribosyl 1-PP | C5H13O14P3 | HMDB0000280 | 16.0 | 389,95 | 388.90 | 177.00 | 25 |
| 5-Phosphoribosyl 1-PP | C5H13O14P3 | HMDB0000280 | 16.0 | 389,95 | 388.90 | 79.00 | 50 |
| 5-Phosphoribosylamine | C5H12NO7P | HMDB0001128 | 1.5 | 229 | 228.0 | 97.0 | 10 |
| 5-Phosphoribosylamine | C5H12NO7P | HMDB0001128 | 1.5 | 229 | 228.0 | 79.0 | 48 |
| Pyruvic acid | C3H4O3 | HMDB0000243 | 15.1 | 88,02 | 87.00 | 43.20 | 4 |
| Ribitol | C5H12O5 | HMDB0000508 | 1.5 | 152,07 | 151.10 | 89.20 | 9 |
| Ribitol | C5H12O5 | HMDB0000508 | 1.5 | 152,07 | 151.10 | 71.10 | 17 |
| Ribitol 5-P | C5H13O8P | n.a. | 13.5 | 232,03 | 231.00 | 97.00 | 12 |
| Ribitol 5-P | C5H13O8P | n.a. | 13.5 | 232,03 | 231.00 | 79.00 | 50 |
| D-Ribose 5-P | C5H11O8P | HMDB0001548 | 13.5 | 230,02 | 229.00 | 97.00 | 10 |
| D-Ribose 5-P | C5H11O8P | HMDB0001548 | 13.5 | 230,02 | 229.00 | 79.00 | 48 |
| D-Ribose 1-P | C5H11O8P | HMDB0001489 | 15.5 | 230,02 | 229.00 | 97.00 | 10 |
| D-Ribose 1-P | C5H11O8P | HMDB0001489 | 15.5 | 230,02 | 229.00 | 79.00 | 48 |
| Ribulose 5-P | C5H11O8P | HMDB0000618 | 14.5 | 230,02 | 229.01 | 97.00 | 14 |
| Ribulose 5-P | C5H11O8P | HMDB0000618 | 14.5 | 230,02 | 229.01 | 79.00 | 48 |
| D-Sedoheptulose 7-P | C7H15O10P | HMDB0001068 | 13.5 | 290,04 | 289.00 | 96.90 | 18 |
| D-Sedoheptulose 7-P | C7H15O10P | HMDB0001068 | 13.5 | 290,04 | 289.00 | 79.00 | 48 |
| D-Sedoheptulose 1,7-bisphosphat | C7H16O13P2 | HMDB0060274 | 21.2 | 370,01 | 369.00 | 97.00 | 14 |
| D-Sedoheptulose 1,7-bisphosphat | C7H16O13P2 | HMDB0060274 | 21.2 | 370,01 | 369.00 | 79.00 | 48 |
| Sorbitol | C6H14O6 | HMDB0000247 | 1.6 | 182 | 181.1 | 101.0 | 13 |
| Sorbitol | C6H14O6 | HMDB0000247 | 1.6 | 182 | 181.1 | 59.0 | 21 |
| Trehalose | C12H22O11 | HMDB0000975 | 1.5 | 342,12 | 341.00 | 89.01 | 12 |
| UDP-GlcNAc | C17H27N3O17P2 | HMDB0000290 | 20.8 | 607,08 | 606.07 | 385.1 | 28 |
| UDP-GlcNAc | C17H27N3O17P2 | HMDB0000290 | 20.8 | 607,08 | 606.07 | 282.2 | 30 |
| Uridine | C9H12N2O6 | HMDB0000296 | 4.0 | 244,07 | 243.1 | 200.00 | 6 |
| Uridine | C9H12N2O6 | HMDB0000296 | 4.0 | 244,07 | 243.1 | 110.00 | 12 |
| Xylitol | C5H12O5 | HMDB0002917 | 1.5 | 152,07 | 151.10 | 89.00 | 8 |
| Xylitol | C5H12O5 | HMDB0002917 | 1.5 | 152,07 | 151.10 | 71.00 | 8 |
| D-Xylose | C5H10O5 | HMDB0000098 | 4.0 | 150,05 | 149.05 | 89.02 | 4 |
| D-Xylose | C5H10O5 | HMDB0000098 | 4.0 | 150,05 | 149.05 | 71.01 | 4 |
| D-Xylulose 5-P | C5H11O8P | HMDB0000868 | 14.3 | 230,02 | 229.00 | 138.98 | 8 |
| D-Xylulose 5-P | C5H11O8P | HMDB0000868 | 14.3 | 230,02 | 229.00 | 79.00 | 36 |
