## Supplemental Table 4 for "Dynamic analysis of sugar metabolism reveals the mechanisms of action of synthetic sugar analogs"

### Supplementary Table S4. Comparative analysis of the relative abundance of nucleotide sugars in 13 organisms or cell lines.

Relative levels of nucleotide sugars in HAP1 cells cultured under different conditions (supplementary Figure 1)

| | CMP-NeuNGc | CDP-ribitol | CMP-NeuNAc | UDP-arabinose | UDP-galactose | UDP-glucose | UDP-xylose | UDP-ManNAc | UDP-GlcNAc | UDP-GalNAc | GDP-mannose | GDP-glucose | UDP-GlcA | GDP-fucose | TDP-glucose | TDP-rhamnose | ADP-glucose | ADP-ribose |
|---|---|---|---|---|---|---|---|---|---|---|---|---|---|---|---|---|---|---|
| t=8 DMEM high | 0,09 | 0,39 | 5,16 | 0,12 | 10,01 | 36,91 | 0,51 | n.d. | 25,56 | 4,40 | 0,74 | 0,00 | 14,60 | 0,80 | 0,01 | n.d. | 0,00 | 0,70 |
| t=8 IMDM high | 0,07 | 0,30 | 3,78 | 0,12 | 10,75 | 40,65 | 0,50 | n.d. | 23,11 | 4,15 | 0,61 | 0,00 | 14,64 | 0,68 | 0,00 | n.d. | 0,01 | 0,63 |
| t=8 DMEM high+ | 0,08 | 0,31 | 5,11 | 0,15 | 10,34 | 37,29 | 0,55 | n.d. | 25,17 | 4,19 | 0,81 | 0,00 | 14,28 | 0,95 | 0,01 | n.d. | 0,00 | 0,75 |
| t=8 DMEM low+ | 0,10 | 0,54 | 4,47 | 0,12 | 10,86 | 42,47 | 0,85 | n.d. | 21,95 | 3,64 | 0,81 | 0,00 | 12,37 | 1,06 | 0,01 | n.d. | 0,01 | 0,76 |
| t=8 DMEM - 2,75 | 0,11 | 0,61 | 4,60 | 0,06 | 10,80 | 39,49 | 0,74 | n.d. | 22,01 | 3,91 | 0,98 | 0,00 | 14,62 | 0,97 | 0,01 | n.d. | 0,01 | 1,07 |
| t=8 DMEM low | 0,07 | 0,71 | 4,98 | 0,08 | 10,07 | 40,01 | 1,02 | n.d. | 22,49 | 3,95 | 0,97 | 0,01 | 13,07 | 1,05 | 0,01 | n.d. | 0,00 | 1,51 |

Relative levels of nucleotide sugars in organisms and cells (Figure 2)

| | CMP-NeuNGc | CDP-ribitol | CMP-NeuNAc | UDP-arabinose | UDP-galactose | UDP-glucose | UDP-xylose | UDP-ManNAc | UDP-GlcNAc | UDP-GalNAc | GDP-mannose | GDP-glucose | UDP-GlcA | GDP-fucose | TDP-glucose | TDP-rhamnose | ADP-glucose | ADP-ribose |
|---|---|---|---|---|---|---|---|---|---|---|---|---|---|---|---|---|---|---|
| Human WT fibroblasts | 0,16 | 1,76 | 4,23 | 0,12 | 6,50 | 20,25 | 0,65 | 0,00 | 34,58 | 22,41 | 1,18 | 0,01 | 6,32 | 1,09 | 0,01 | 0,00 | 0,01 | 0,73 |
| HEK293 | 0,23 | 0,51 | 9,11 | 0,09 | 6,47 | 19,53 | 0,47 | 0,00 | 32,46 | 21,26 | 1,86 | 0,00 | 6,15 | 1,45 | 0,00 | 0,00 | 0,00 | 0,40 |
| HAP1 | 0,04 | 0,32 | 3,14 | 0,09 | 6,47 | 22,95 | 0,28 | 0,00 | 31,74 | 20,85 | 1,27 | 0,01 | 11,87 | 0,70 | 0,01 | 0,00 | 0,00 | 0,25 |
| Mouse C2C12 myoblasts | 0,25 | 2,34 | 5,94 | 0,50 | 14,02 | 41,69 | 1,47 | 0,00 | 10,73 | 7,31 | 0,75 | 0,05 | 13,38 | 0,86 | 0,04 | 0,00 | 0,03 | 0,64 |
| Mouse C2C12 myotubes | 0,82 | 7,52 | 3,87 | 0,02 | 8,64 | 21,06 | 0,67 | 0,00 | 28,27 | 17,96 | 0,84 | 0,00 | 9,71 | 0,56 | 0,00 | 0,00 | 0,01 | 0,04 |
| Zebrafish adult | 4,89 | 1,63 | 27,16 | 0,34 | 0,80 | 2,67 | 0,47 | 0,00 | 35,73 | 23,14 | 0,17 | 0,02 | 2,22 | 0,53 | 0,02 | 0,08 | 0,04 | 0,09 |
| Zebrafish embryo | 4,25 | 1,55 | 12,70 | 0,22 | 3,72 | 12,90 | 0,58 | 0,00 | 35,99 | 23,17 | 0,08 | 0,02 | 1,83 | 0,86 | 0,01 | 0,00 | 0,02 | 2,11 |
| Canton R male flies | 0,00 | 0,00 | 0,03 | 0,32 | 14,08 | 45,98 | 0,24 | 0,00 | 22,13 | 4,64 | 5,26 | 0,08 | 3,49 | 0,57 | 0,04 | 0,00 | 0,17 | 2,97 |
| Canton R female flies | 0,00 | 0,01 | 0,03 | 0,26 | 12,62 | 37,98 | 0,24 | 0,00 | 27,79 | 5,55 | 7,24 | 0,06 | 4,53 | 0,65 | 0,04 | 0,00 | 0,15 | 2,86 |
| Canton R Virgin females | 0,00 | 0,01 | 0,03 | 0,41 | 13,25 | 28,90 | 0,27 | 0,00 | 34,73 | 6,73 | 4,80 | 0,08 | 3,71 | 0,51 | 0,07 | 0,00 | 0,21 | 6,28 |
| Yeast | 0,00 | 0,01 | 0,00 | 0,00 | 2,31 | 46,24 | 0,00 | 0,00 | 22,41 | 0,04 | 25,98 | 0,04 | 0,02 | 0,00 | 0,01 | 0,00 | 0,04 | 2,91 |
| gram positive bacteria | 0,02 | 0,05 | 0,04 | 0,01 | 4,58 | 19,57 | 0,02 | 0,00 | 33,85 | 3,58 | 0,03 | 0,02 | 0,01 | 0,03 | 0,83 | 35,63 | 0,16 | 1,57 |
| Bacteria gram negative | 0,00 | 13,97 | 0,00 | 0,02 | 3,02 | 10,51 | 0,01 | 0,41 | 28,85 | 18,54 | 0,07 | 0,01 | 0,03 | 0,00 | 0,06 | 22,81 | 1,22 | 0,46 |
