## Supplemental Table 5 for "Dynamic analysis of sugar metabolism reveals the mechanisms of action of synthetic sugar analogs"

**Table S5. Overview of the effect of ManNPoc and SiaNPoc labeling in fibroblasts on all nucleotide sugars and incorporation of the Poc group.**

The relative abundances of nucleotide sugars are shown after incubation of fibroblasts with Ac5ManNPoc, Ac5NeuNPoc, Ac5NeuNAc or PBS. In addition, the incorporation of ManNPoc and SiaNPoc in nucleotide sugars is shown as ratio of the theoretically produced Poc derivatized nucleotide-sugar versus endogenous, non-modified nucleotide sugars.

| Sample | CDP-ribitol | CMP-NeuNAc | UDP-galactose | UDP-glucose | UDP-arabinose | UDP-xylose | UDP-GalNAc | UDP-GlcNAc | GDP-glucose | GDP-mannose | UDP-GlcA | CMP-NeuNPoc | GDP-fucose | UDP-HexNPoc | TDP-glucose | ADP-glucose | ADP-ribose |
|---|---|---|---|---|---|---|---|---|---|---|---|---|---|---|---|---|---|
| **Average** | | | | | | | | | | | | | | | | | |
| ManNPoc | 0,75 | 4,20 | 11,24 | 40,16 | 2,14 | 2,25 | 3,67 | 21,90 | 0,08 | 0,33 | 9,40 | 0,75 | 1,44 | 0,16 | 0,30 | 0,11 | 0,90 |
| NeuNPoc | 0,75 | 4,32 | 11,38 | 36,50 | 1,99 | 2,08 | 4,58 | 23,97 | 0,09 | 0,32 | 9,83 | 1,40 | 1,34 | 0,00 | 0,28 | 0,11 | 0,85 |
| NeuNAc | 0,77 | 5,38 | 13,10 | 38,10 | 1,49 | 1,56 | 3,81 | 23,47 | 0,04 | 0,33 | 9,87 | 0,00 | 1,22 | 0,00 | 0,16 | 0,05 | 0,45 |
| Control (PBS) | 0,78 | 4,91 | 11,83 | 38,22 | 1,77 | 1,86 | 4,29 | 22,88 | 0,06 | 0,37 | 10,19 | 0,00 | 1,24 | 0,00 | 0,23 | 0,10 | 0,71 |
| **Percentage Poc labeling** | | | | | | | | | | | | | | | | | |
| ManNPoc | n.d | 18 | n.d | n.d | n.d | n.d | n.d | 0,6 | n.d | n.d | n.d | n.d | n.d | n.d | n.d | n.d | n.d |
| NeuNPoc | n.d | 32 | n.d | n.d | n.d | n.d | n.d | n.d | n.d | n.d | n.d | n.d | n.d | n.d | n.d | n.d | n.d |
| NeuNAc | n.d | n.d | n.d | n.d | n.d | n.d | n.d | n.d | n.d | n.d | n.d | n.d | n.d | n.d | n.d | n.d | n.d |
