## Supplemental Table 6 for "Dynamic analysis of sugar metabolism reveals the mechanisms of action of synthetic sugar analogs"

### Table S6. Overview of the effects of 3Fax-NeuNAc and NeuNAc in B16-F10 cells on all nucleotide sugars.

The relative abundances of nucleotide sugars are shown after 48 h incubation of B16-F10 cells with Ac5-3Fax-NeuNAc, Ac5NeuNAc or PBS. In addition, the incorporation of F-Sia is shown as ratio of the theoretically produced fluoro derivatives versus endogenous, non-modified nucleotide sugars.

| Sample | CDP-ribitol | CMP-NeuNAc | CMP-3Fax-NeuNAc | UDP-galactose | UDP-glucose | UDP-arabinose | UDP-xylose | UDP-GlcNAc | UDP-GalNAc | GDP-glucose | GDP-mannose | UDP-GlcA | GDP-fucose | ADP-glucose | TDP-glucose | ADP-ribose |
|---|---|---|---|---|---|---|---|---|---|---|---|---|---|---|---|---|
| **Average** | | | | | | | | | | | | | | | | |
| 3Fax-NeuNAc | 0,231 | 0,291 | 34,474 | 11,153 | 33,117 | 0,086 | 0,680 | 1,167 | 15,939 | 0,000 | 0,033 | 2,068 | 0,618 | 0,014 | 0,001 | 0,101 |
| NeuNAc | 0,331 | 9,564 | 0,006 | 15,937 | 48,493 | 0,124 | 1,040 | 1,407 | 19,111 | 0,001 | 0,051 | 2,790 | 0,932 | 0,014 | 0,001 | 0,153 |
| Control (PBS) | 0,367 | 8,278 | 0,034 | 16,096 | 49,014 | 0,122 | 1,037 | 1,519 | 19,640 | 0,001 | 0,049 | 2,738 | 0,895 | 0,016 | 0,006 | 0,152 |
